# Mineral geochemistry and mycorrhizal allocation define root architectural strategy during early vascular plant colonization

**DOI:** 10.64898/2026.07.06.736658

**Authors:** DG Zaharescu, Jennifer K. Presler, Miranda Galey, Alexandra Salinas, Kexin Li, Carmen I Burghelea, Estefanía C. Roldán-Nicolau, Valentina Vavages, Katerina Dontsova, Raina M. Maier, Jon Chorover, Rebecca Lybrand, Travis Huxman

## Abstract

The emergence of vascular plants on land is one of evolution’s greatest triumphs. This success was contingent on the capacity of roots and their symbionts to acquire resources from exposed geology. However, how rock chemistry shapes plant root architectural strategies, and their return on investment during early ecosystem colonization remains poorly understood. Here we use a two-year mesocosm experiment with *Bouteloua dactyloides* grass and an arbuscular mycorrhizal symbiont, grown on four mineral substrates of contrasting composition, to show that rock geochemistry predictably determines root topological strategy, from herringbone architecture on nutrient-poor granite to dichotomous-like branching on nutrient-rich basalt. Substrate identity governed investment allocation between root complexity and biomass, with plants consolidating existing transport pathways as weathering-derived nutrients subsided. Traits associated with exploratory effort were generally decoupled from those related to biomass buildup. In **basalt** and **rhyolite** plants preferentially invested in complexity, generating the largest numbers of prospective tips for mining and biomass buildup; in **granite**, plants chose a surviving strategy, limiting branching to preserve biomass; while in **schist**, plants balanced biomass with complexity, extending growth on low investment, which increased tissue density. Surprisingly, mycorrhizal fungi did not alter the whole root system size, but reallocated investment between specific root orders, discouraging investment in embryonic roots in some substrates, and stimulating lateral expansion of the rooting system in others. This extends the functional balance mechanism from plant to the plant–fungus system.

The extensive phenotypic plasticity of the root-mycorrhiza system shown here provides an evolutionary space for natural selection, which must have played a crucial role in the success of plants on land in the past, and is crucial for understanding plant ecological dynamics today.

## 1. Introduction

Vascular plants are among Earth’s most successful land colonizers, and architects of the terrestrial biosphere. They grow in fresh and saline waters, occupy deserts, create forests, inhabit mountain tops, and through their above and belowground branching, they create habitats that nourish a great diversity of life forms. Such evolutionary success was made possible almost exclusively due to the capacity of vascular plants to germinate, acquire energy, and grow within rather narrow environmental boundaries in which seeds settle. Among major plant organs, roots have evolved various acquisitive (absorptive) and non-acquisitive (structural, storage, production of exudates, foraging and other secondary functions) compartments (Klimešová et al., 2018) that allowed plants to efficiently strategize energy use in ways that maintained fitness across a wide diversity of climate and ecosystem constrains. Roots have therefore been described as multidimensional systems, with groups of traits often capable of responding independently to resource variation in the environment (Kramer-Walter et al., 2016).

The root phenotypic plasticity – the capacity to change morphology to cope with stress, is one of the best tools used by plants to maximize energy intake while adapting to resource fluctuations in the substrate. The ability to explore water and nutrient hotspots, avoid toxic chemicals, unfavorable temperature and solar radiation are few of the strategies the root system can use during development. Similarly, developing roots connect to pre-existing networks of fungi and microbes, which expand the rhizosphere, thus a plant’s reach in the heterogeneous nutrient space (Hinsinger, 1998; Solís-Domínguez et al., 2011; Zhang et al., 2024; Cao et al., 2024). By occupying niches an order of magnitude smaller than roots, fungal hyphae and microbes have a larger area of interaction with the substrate, and more access to nutrients.

Similar to other anatomical and physiological strategies shared by life, such as symmetry, allometry (the proportional development of a body plan), photosynthesis, and respiration/fermentation, several energy investment strategies allow plants to maximize energy return (e.g. biomass) on energy invested, improving fitness in a competitive environment. Two of the most relevant are: **(i)** maximizing the energy (biomass) returned per functional structures (biomass-producing units, e.g. ribosomes) built, and **(ii)** maximizing the energy returned per work performed by such structures (Burnetti et al., 2026). The first case is a low energy investment strategy suitable for situations of high growth/reproduction rate under abundant resources that allow structures to be replicated in large numbers, and corresponds to the fast end-member of the fast-slow growth continuum in root traits outlined by Chapin, 1980, Freschet et al., 2010; Bergmann et al., 2020; Klimešová and Herben, 2021, and others. It is commonly used by highly competitive generalists that grow quickly to take advantage of abundant resources as early as possible, e.g. annual grasses, first herbaceous colonizers of an exposed soil (Chapin, 1980). The second case requires large energy investment to increase nutrient acquisition and store in otherwise nutrient-limited environments, and assure good fitness and competitive success. The latter is used by specialists that invest more energy over longer periods of time (e.g. perennial grasses and woody plants), and represent the slow end-member of the fast-slow growth continuum in root traits (Bergmann et al. 2020, Klimešová and Herben 2021).

In physiological terms, the allocation of costly metabolic enzymes to efficiently process energy resources acquired from the environment leads to a compromise between investment in metabolic output and growth rate; with slow growth and comparatively high metabolic output in scarce resources, due to less resource allocation to new enzyme producers (i.e., N and P to ribosomes, cells). And, rapid growth and low metabolic output occurring in abundant resources due to more investment in new structures. This process has been clearly exemplified in microbes (Wortel et al., 2018; Burnetti et al., 2021).

In allometric terms, for a given amount of available energy resource in an organism, the more energy is allocated to a developing trait, the less is available for developing other traits. Strategizing the energy investment in development of different body parts is critical for organism survival during stress, and has been demonstrated in plants, invertebrates and vertebrates (Stern and Emlen, 1999; Weiner, 2004).

Two major root branching strategies have been used by vascular plants throughout their evolution to maximize return on investment: **(i)** *herringbone strategy*, a coarse - scale foraging strategy for extensive soil exploration with limited number of lateral roots (primarily confined to main embryonic axes). This strategy is efficient at acquiring mobile nutrients (e.g. nitrate) in bulk solution, and involves significant photosynthetic carbon cost in building up the root system (Fitter, 1985, Fitter and Stickland, 1991; Beidler et al., 2015). And **(ii)** *dichotomous strategy*, a fine-scale intensive foraging strategy with extensive branching for acquiring diffusion-restricted resources such as Fe and P that quickly immobilize as oxides (Fitter and Stickland, 1991).

Since energy allocation strategies play a crucial role in an organism/population survival, they could have evolved with the raise of multicellularity. To what extent have such strategies played out when vascular plants diversified on land in early Phanerozoic is not known; however, in ecological terms, energy tradeoffs within individuals, and between individuals and species allowed species to efficiently occupy new niches, contributing to niche stratification and ecosystem spread-out (Wilkinson, 2023). Today, such strategies help in species dispersal, e.g. on land exposed by volcanic eruption, after glaciers retreat, as well as during the upward and pole-ward migration of populations with climatic changes (Anthelme et al., 2021). Grasses are one of plant evolution’s most recent and successful branch (at least 60ma; Prasad et al., 2005), which currently cover about 40% of land surface, and 69% of the Earth’s agricultural land area (Bardgett et al., 2021). The efficient energy allocation strategies evolved by grasses gave rise to the successful spread of C4 type photosynthesis (Osborne and Beerling, 2005), and have contributed to their widespread dominance in various ecosystems.

The evolution of a plant’s root system has also benefited from an extensive microecosystem of fungi and microbes in the soil environment. For instance, fungal mycorrhization coincided with the development of primordial roots during the land colonization by plants, from early Triassic (ca. 250 million years ago) for ectomycorrhiza (Sato, 2023) to more than 400 million years ago for arbuscular mycorrhiza (Remy et al., 1994; Brundrett, 2002). Mycorrhizae are also integral to the modern below-ground acquisitive compartment in plants (Klimešová et al., 2018); hence, their critical role in root development strategies and plant success cannot be neglected. Because root development is shaped by the most rewarding return on investment strategy, which in natural ecosystem is aided by the surrounding microbiome, its effect could be captured in its allometric patterns of growth.

Mineral substrates under early weathering are perhaps the best analogs of new land niches today, and in the past when vascular plants diversified on the rocky land. Due to nutrient availability strictly related to their mineral composition, such substrates could offer a clear window into how roots adapt to different organic-limited environments during resource exploration. Our overarching goal was to better understand how root exploration and exploitation strategies in vascular plants develop during germination and growth in response to bedrock mineralogy. We conducted a two-year mezocosm-scale experiment, where we examined how the vascular plant *Bouteloua dactyloides* (Buffalo grass) interacted with four granular substrates (basalt, rhyolite, granite and schist) in both, the presence and absence of arbuscular mycorrhiza *Rhizophagus irregularis (*previously known as *Glomus intraradices)*.

### We hypothesized that

**(i)** Root architecture measures capture different strategies plants use during development; **(ii)** Plants employ different root geometries to maximize fitness in different mineral substrates, and such strategies will be reflected in the above and below-ground organs; and **(3)** fungal symbiosis will influence the root system energy investment tradeoff. A major advantage of using granular rock is that the substrate reduces the inherent complexity of developed soil environment, while magnifying the response of the root system to specific geological settings.

## 2. Results and discussion

### 3.1 Global descriptors of plant development

Of the four rock types, rhyolite supported, on average, the highest survival rate of pre-germinated seeds, followed by basalt, granite, and schist (Fig. s3A). Substrates inoculation with arbuscular mycorrhiza coincided with increased plant survival rate in all rocks except basalt, where survivability was lower (Fig. s3B). Increased root longevity under mycorrhiza associations has been reported for infertile habitats in early studies (Mosse, 1973; Chapin, 1980), outlining the important role of fungal associations to plant survival in low-nutrient settings. After an initial drop due to seedling die-offs, survival rates did not change with time in any of the rocks, evidence of plant adaption to un-amended mineral substrate (as the sole nutrient source) and milliQ water (Fig. s3C). Total biomass, measured across the four sacrifice time points (132, 252, 465 and 584 days) was lower in the shoots compared to the roots, indicating a strong energy investment in the belowground nutrient-acquisition organ (Fig. s3D). While schist hosted the lowest total plant biomass among rocks, the root mass fraction (root/plant) was also largest in schist (Fig. s3E), which associated to increased root tissue density (largest among rock, Fig. s3F). High root : shoot biomass values are common in infertile habitats across species, as a phenotypic response to reduced nutrient availability (Chapin, 1980; Dennis and Johnson, 1970; Christie and Moorby, 1975). Root density and branching have also been negatively related to growth rate and soil fertility in tree species, in unison with other above-ground traits (Kramer-Walter et al., 2016), altogether implying that the schist in our system was among the least fertile substrates for *B. dactyloides*.

### 3.2 Root investment strategies

Plants have a limited pool of internal resources to help maximize external nutrients/energy intake during development, and we hypothesize that a highly variable but restricting substrate affects patterns of allometric growth. An initial Principal Component Analysis grouped the 18 measured variables into two principal components (PCs) that represent major root traits and their functions, which together accounted for 80.24% of total variance in root morphological characteristics (Fig. 1A, B). PC1, which we interpret as root complexity, explained 40.3% of total variance and represents variables that can be ascribed to root investment in exploratory effort (i.e., count variables including total number of axes, number of lateral and tertiary axes, links, root branching, and tips. Altitude (largest nutrient path by segment number), external path length (number of segments from root base leading to tips) and the total length of tertiary roots are also associated with PC1. PC2, interpreted as root biomass investment, explained the ensuing 39.9% of total variance (Fig. 1A, B), and included grouped traits that are associated with root biomass buildup effort. Surface area, representing a plant’s nutrient-extraction interface, which directly responds to a plant’s physiological needs, leads the PC1 with the highest loading. Root thickness (average diameter), volume, and length were also positively associated with surface area. PC2 grouped root segment variables included diameter (link diameter), surface area (link SA), length (link L), as well as developmental variables including main and lateral root lengths. Patterns of biomass-related variables independent of morphological variables were also observed in a variety of woody plants (Yang et al., 2021), suggesting a decoupled root complexity - biomass across the vascular plants domain.

**Fig. 1.**
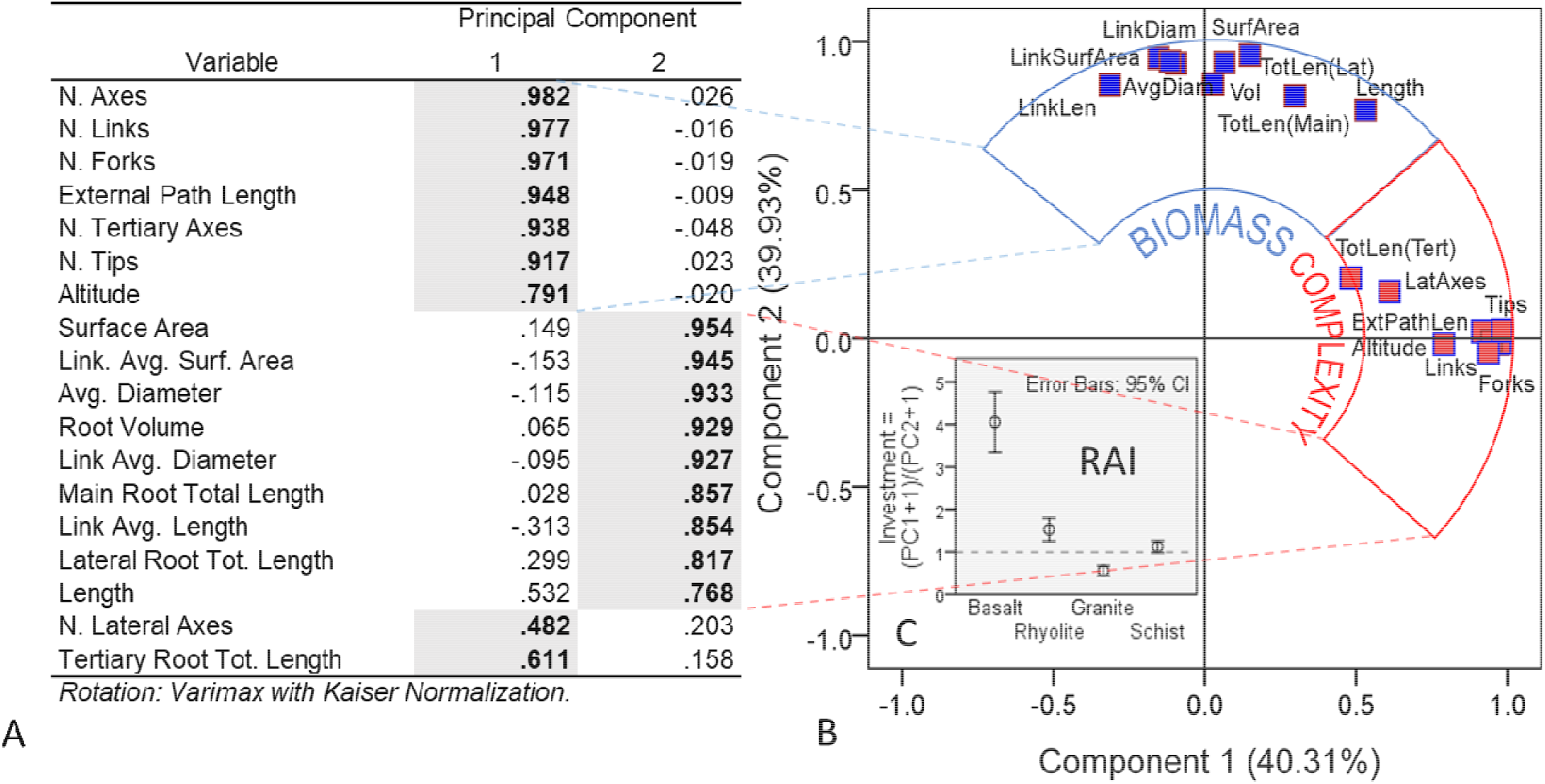
Global descriptors of root development. (A, B) Principal Component Analysis grouping the morphological traits of *B. dactyloides* root, based on their similarities. (C) Root Allometric Investment Index (RAI), represented by the ratio between regression factor scores of PC1 and PC2 denoting investment in allometric growth (complexity vs. biomass) of the root system in the various rock substrates. N = 780 plants.

Allometric effort, estimated by Root Allometic Index (RAI) differed among substrates when comparing investment in complexity to investment in biomass (Fig. 1C). Plants in basalt and rhyolite showed preferential allometric investment in root complexity (values above unity), with plants in basalt having by far the largest RAI values (about four times larger than in rhyolite; Fig. 1C). Plants in granite, on the other hand, invested resources mostly in root biomass (values below unity), while plants in schist invested equally in developing root complexity and biomass. The differences in allometric growth among substrates were significant, meaning that rock type dictated how *B. dactyloides* strategized root development. Presence of fungal mycorrhiza, on the other hand did not significantly affect RAI as indicated by similar RAI values between inoculated and non-inoculated plants, showing limitations of mycorrhizal effect on plant energy allocation, particularly during growth on mineral substrates.

### 3.3 Investment in Root Complexity and its Indicators

Our calculated Altitude Topological Index (ATI), which codifies the proportional relationship between exploratory growth phases (altitude) and absorptive segments generation (tips number) into architecture strategies (*Fitter, 1987; Beidler et al., 2015*), ranged from ATI = 0 (dichotomous) to ATI=1 (herringbone), with most roots plotting within the herringbone domain (Fig. 2A). The two strategies allow fine roots to either maximize resource acquisition by exploring and mining a certain volume more intensely (dichotomous), or become more extensive in their search for energy resources in volumes with more diffused nutrients (herringbone; Löhmus et al., 2006). There were significant differences among the four rock substrates, with basalt supporting the most branched roots, and schist and granite, the least (Fig. 2A, B). This means that plants in basalt explored the available soil volume more thoroughly and increasing mycorrhizal tip proliferation (lowest ATI), while plants in granite grew more extensive roots to allow them to prospect further. A similarly wide breadth of branching strategies for a single species was also found in *Arabidopsis* root systems under different nutrient supplies (López-Bucio et al., 2003), and *Pinus taeda*, a gymnosperm older on the evolutionary tree, with plants in environments with less access to nutrients preferring a herringbone investment strategy (Beidler et al., 2015). Differences in branching strategies also suggests that such large phenotypic plasticity within species in terrestrial plants is likely an early evolutionary trait that has helped vascular plants cope with highly variable substrate conditions and increasingly competitive ecosystem environments.

**Fig. 2.**
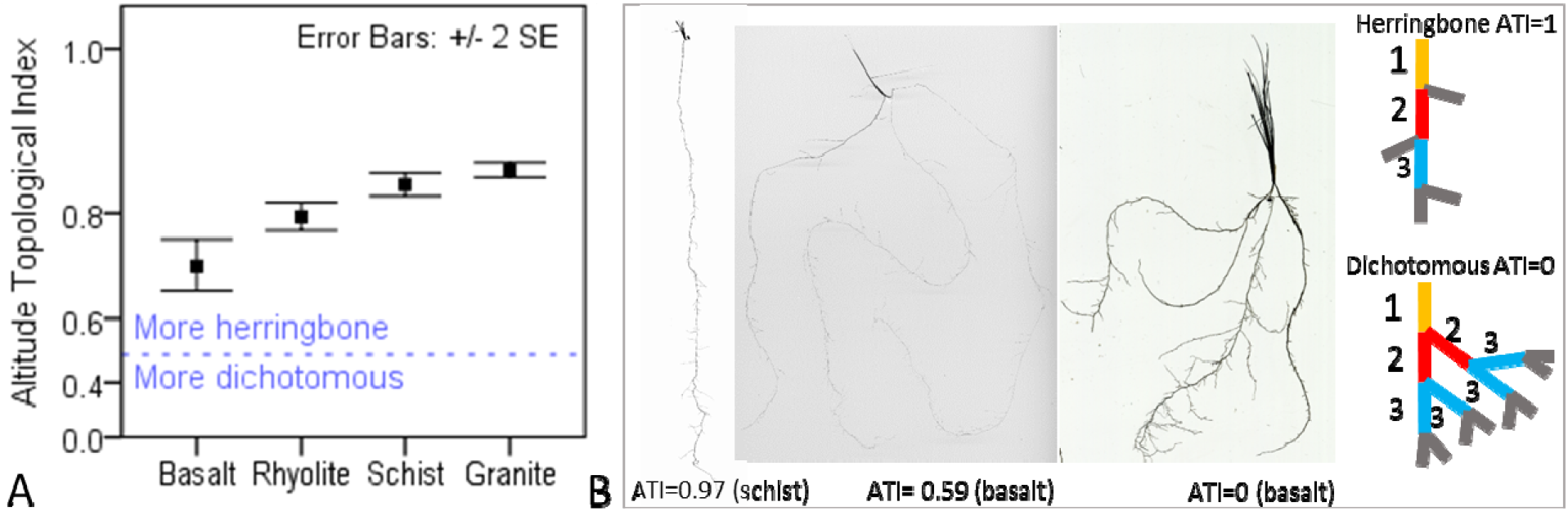
Global topology. (A) Root branching pattern preference in *B. dactyloides* grown in four rock substrates as described by Altitude Topological Index (ATI; Fitter, 1987); N = 644 plants. (B) Examples of herringbone and dichotomous topologies end members, together with their structures. Colors represent sequence of root links/segments formed during the first (1, orange) second (2, red) and third (3, blue) root growth phases, together with terminal/external segments (gray).

Surprisingly, during development plants in rhyolite grew the smallest maximum path for nutrients through the root system by segment number (quantified as *altitude* - an ATI component), while largest were in schist and other rocks (mean = 74.9 segments in rhyolite, mean = 86.9 segments in schist; Fig. s4A). Most sequential development occurred in the first 6 months, when mineral weathering was also stronger (Zaharescu et al., 2019), with mycorrhiza presence significantly contributing to the path shortening in rhyolite (Fig. s4B, C). This is further evidence of various substrate-dependent nutrient transport strategies through the root system used by a single plant species and its fungal symbiont. There was an increase in root system density (approximated by *external path length*) over time across rocks, with significant differences emerging toward the second part of the growth period (Fig. s4E), when nutrient leaching also decreased (Zaharescu et al., 2019). Granite plants had about 30% smaller root system density (Fig. s4D), which shows a weak development. Studies using nutrient enrichment have shown dependence of external path length in woody plants to CO_2_ and N fertilization/ mineralization, with increased external path length under enriched CO_2_ atmosphere (Beidler et al., 2015), although this mechanism could not be the case in our study.

Development of new roots is initiated through cell division in the apical and lateral meristems, and the generation of tips, which are important in functional root branching. The largest number of tips was generated by plants in rhyolite and basalt (Fig. 3A). As expected, fine root class generally had the largest number of tips (Fig. 3B). Basalt and schist supported the largest tips density per plant for the thinnest roots (0-0.3mm thickness), with widest spread in basalt, suggesting greater fine tips variability (Fig. 3B, C). Rhyolite presented the largest tips number across thickness classes, and stimulated the development of slightly thicker fine roots than the other rocks. Granite showed a higher frequency at lower tip counts suggesting the predominance of plants with fewer fine tips (Fig. 3C). Mycorrhizal inoculation coincided with less tips density in fine to medium-thickness roots in rhyolite, and in the finest root class in granite, showing an early fungal influence on plant investment in new structures (Fig. s5).

**Fig. 3.**
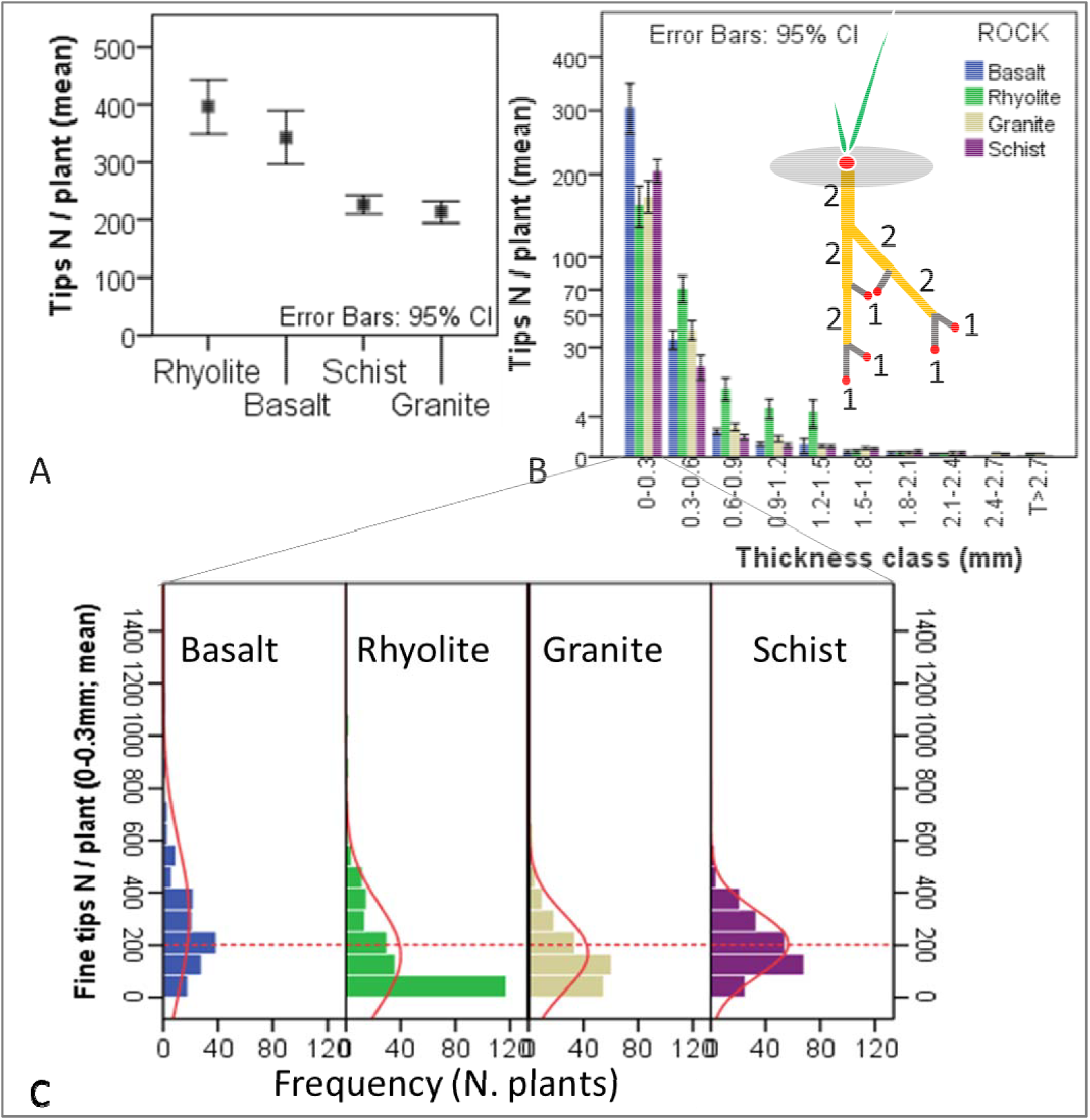
Root tips. (A) Effect of rock type on tips abundance in *B. dactyloides* over 2 years growth. (B) Tips abundance for each root thickness class across rocks. Inset diagram illustrates tip-bearing absorptive (1) and transport (2) segments, with apical growth meristems highlighted in red. (C) Fine tips (0 – 0.3mm diameter) frequency distribution across rocks. Superimposed red lines provide smoothed representations of the distribution shapes, with dashed horizontal line set at group average. N = 780 plants.

Functionally, root tips represent a plant’s investment in generating nutrient prospection and uptake pathways, which ultimately determines the complexity of root geometry in relation to nutrient source and abundance (exploration efficiency), and plant needs. Tips lead the emergence of root hair regions - the main nutrient uptake structures (70 - 90% of total root area; Holz et al., 2017), and rhizosphere microhabitats in areas where nutrients can be accessed. Tips grow in response to plant allocating growth hormones (e.g., auxins, cytokinins, ethylene), to local epidermic regions (meristems) that sense differences in nutrient concentrations between root and the pore environment in a recurring, memory-imprinting pattern upregulated by light/sugar production (Lopez-Bucio et al., 2003; Reyes-Hernández and Maizel, 2023). This environment-root feedback mechanism is upregulated by melatonin - an ancient enzyme common to unicellular and multicellular eukaryotes that activates genes related to the production of auxin, and root-tip meristem initiation, as well as genes involved in nitrate and Zn transport (Liang et al., 2017). It has been shown that the melatonin-auxin signaling pathway orchestrates post-embryonic root architecture by inhibiting the growth of pre-existing (embryonic) roots, and stimulates lateral root and root hairs formation and shoot growth in a variety of grasses (Hernandez-Ruiz et al., 2005; Overvoorde et al., 2010; Zhang et al., 2014). Other hormones, such as gibberellic acid, can also balance below ground tip production with above ground shade response (van Gelderen et al., 2023); but the comparatively abundant tip generation in basalt and rhyolite compared to other rocks in our study is most likely due to the geochemical-embryological feedback in a generally nutrient-limited environment. Mycorrhiza’s increased access to chemical energy resources and its acquisition efficiency could also explain why root tip generation decreased in plants inoculated with the fungus, as plants could divest limited energy/nutrients from building roots to other functions. This supports the view that mycorrhizal fungi are an extension of the root system (Bunn et al., 2024). The observed differences in root tip production may also lead to modifications in other root architecture parameters such as links and axes.

Episodic tip production initiates lateral branch development, which establishes distinct nutrient transport pathways (quantified as root axes) that connect fine acquisitive root segments to aerial stems for resource translocation (Fig. 4A). Roots in basalt experienced more branching episodes (higher total number of axes) compared to the other substrates, indicating an abundant and more complex root system (Fig. 4B inset). Lateral (secondary) and tertiary axes were also more numerous in basalt, reflecting more energy invested per biomass in exploratory activity and nutrient uptake than in other rocks (Fig. 4 C, D inset). The basalt scoria in our study is comparatively more abundant in Mg, Fe, K and bioavailable P (elements critical to photosynthesis and plant growth), and presents higher solubility due to its vesicular morphology and glass content (Burghelea et al., 2015; Zaharescu et al., 2019), which could explain their increased exploratory investment. Surprisingly, our results also suggest that in a mineral environment generally deprived of easily available P and N, and no expected nutrient gradient along the column, the growth energy provided by photosynthesis-derived C and more abundant major cations (i.e., Mg, Fe, K) in basalt was redirected from main (embryonic) root transport pathways towards developing lateral roots to improve P harvest. Conversely, granite provided the lowest water-available P (Zaharescu et al., 2019), which coincided with the plants developing the smallest number of branching and a weak biomass buildup (Fig. 4B inset, Fig. s3D).

**Fig. 4.**
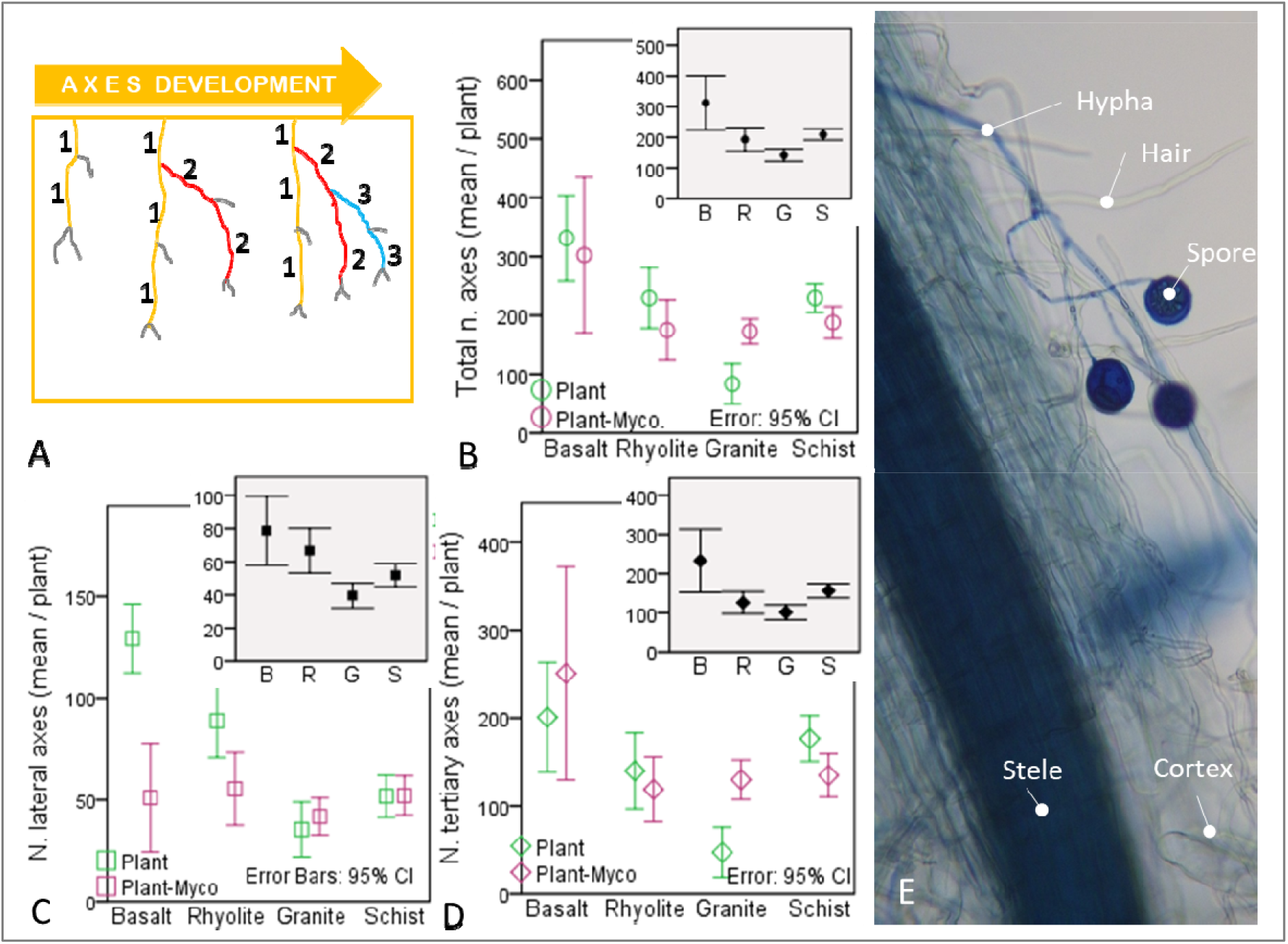
Sequential root development. (A) Illustration of branching episodes and the organization of links/segments into (1, orange) primary, (2, red) secondary, and (3, blue) tertiary axes, together with external links (gray). (B) Total number of axes, (C) lateral axes, and (D) tertiary axes in roots inoculated and non-inoculated with arbuscular mycorrhiza (*R. irregularis*) across rock substrates (pooled across biotic treatments in inset plots), after two years of growth. N= 405 plants; B - basalt, R – rhyolite, G – granite, and S – schist. (E) Stained root highlighting fungal hyphae and spores from plants in basalt after 130 days.

Mycorrhiza inoculation had an inhibitory effect on the number of secondary axes in basalt and rhyolite, which also mirrored the overall tips number in rhyolite (Fig. 4C). The extensive branching of the fungal symbiont, together with their small diameter (around 2µm, or 10- to 100-fold smaller than that of roots) and unique adaptation to locate and mine nutrient-rich mineral sites (particularly P), greatly expand nutrient access pathways, and the transport geometry that roots alone cannot achieve, effectively acting as a root system extension (Bunn et al., 2024). We speculate that the presence of fungi decreased the plants need to invest limited growth resources into root branching, as the energy required for supporting hyphae is likely lower than that needed to produce an equivalent surface area of roots (of much greater volume). In granite (substrate that also supported the lowest tertiary roots count, and low available P), tertiary branching episodes increased under mycorrhiza treatment (Fig. 4B, F).

The greater weight of root axes on the overall geometry parameters in the PCA compared to other parameters (i.e. highest loading value in PC1; Fig. 1A) suggests that from an energy investment standpoint branching takes priority over other root characteristics, as they could represent a more direct plant feedback to depleting chemical energy in the environment.

*Root forks*, a correlate of branching episodes and tips proliferation in PC1 (Fig. 1A, B) were significantly more abundant in rhyolite rock (Fig. 5A inset), a substrate that also supported the largest column plant biomass and seedling survival (Fig. s3A, D). Forks generation (i.e. branching) initiates the organization of complex relationships between plant and the heterogeneous soil environment, including increasing soil exploration volume, providing nutrient and water capture and redundant vascular transport pathways, and anchorage strength (Hodge, 2004; Lynch, 1995, 2019; Pierret et al., 2007; Ennos, 2000). By increasing nutrient interception probability (greater substrate volume exploration), plants in rhyolite seem to successfully invest limited C resources in branching to adapt to nutrient restriction stress and improve biomass buildup. Such pronounced investment in root architectural complexity represents an adaptive response to rhyolite, as proliferation of lateral roots is triggered by regions of high availability of limiting nutrients, combined with a decrease in allocation to other parts of the root system (Chapin, 1980; Lynch, 2019). An enhanced branching facilitates efficient nutrient foraging relative to root construction costs, and it is an expression of optimal resource allocation theory (Bloom et al., 1985). Notably, rhyolite plants also exhibited the most substantial temporal decline in fork density, a pattern also evident, though more attenuated, across the other substrates (Fig. 5A). This progressive reduction in lateral root initiation reflects diminishing biomass buildup most likely due to declining nutrient availability following the initial weathering-driven nutrient pulse as previously shown in this experiment (Zaharescu et al., 2019).

**Fig. 5.**
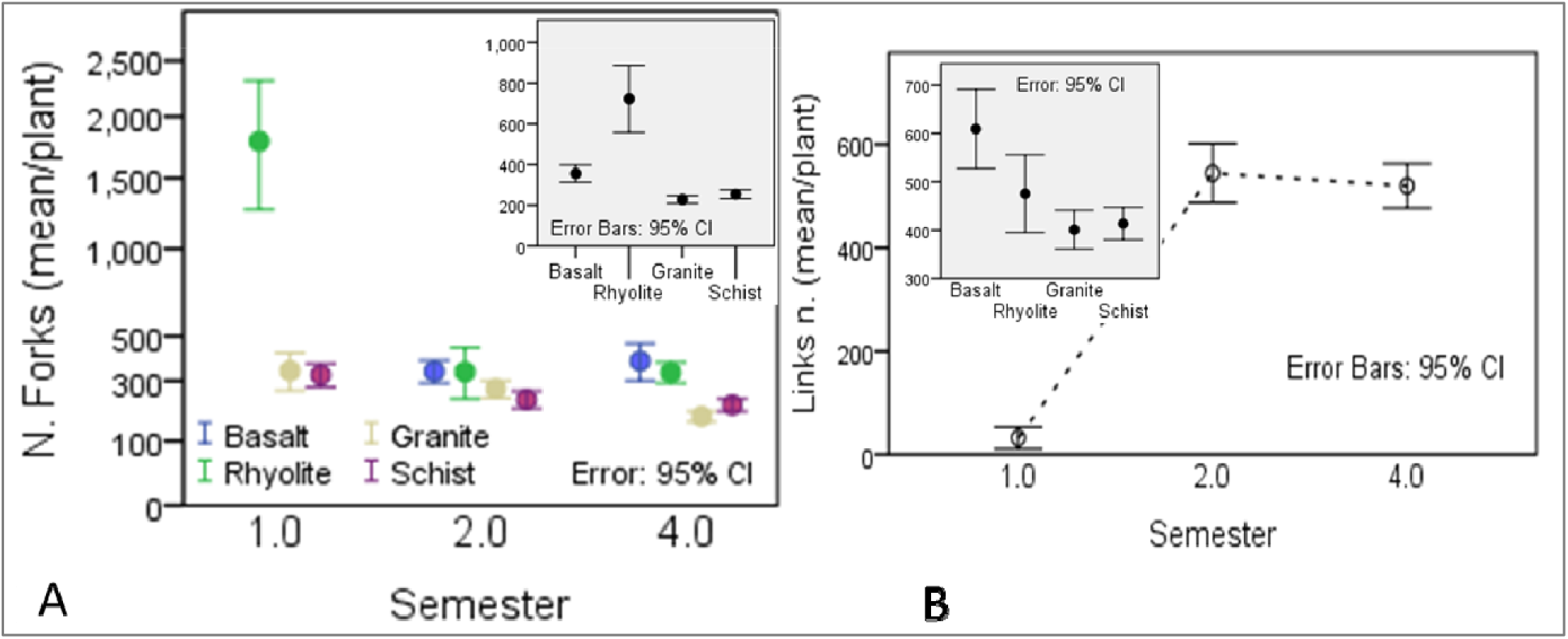
Root branching and segmentation. Effect of time (A), and rock (inset) on the overall development of roots forks in Bouteloa dactyloides over 2 years experiment. (B) Root links number generation along time and across rocks (inset). N=780 plants.

*Root segmentation* (measured by links count), which controls the direction and rate of root growth and determines root functional capacity, was larger in basalt (Fig. 5B inset). Similar to forks, links number increased across all substrates during the first 6 months of growth before stabilizing (Fig. 5B). This pattern suggests that plants in basalt redirected growth resources toward enhanced segmentation – a strategy aligned with dichotomous branching architecture (Fitter and Stickland, 1991) – thereby simultaneously producing lateral roots and enhancing root system connectivity. Such architectural modifications enhance a plant’s volumetric exploration ability and enable flexible resource reallocation when needed (Fitter, 1987; Lynch, 1995). This response to substrate limitation in basalt parallels nutrient-induced architectural plasticity documented in herbaceous species. Experimental manipulations in *Arabidopsis thaliana* showed that P, N, and S deficiencies increased root segmentation (Williamson et al., 2001; Lopez-Bucio et al., 2003; Gruber et al., 2013). Woody species also exhibit similar responses despite their contrasting tissue stoichiometry. For instance, a field study on 14 year-old *Pinus taeda* (of lower N content than herbaceous plants) growing in an experimental forest, revealed decreased root segmentation following nutrient addition, though elevated CO_2_ reversed this trend (Beidler et al., 2015). The convergence of these responses across herbaceous and woody lineages suggests that trait-substrate feedbacks governing belowground resource exploration represent evolutionarily conserved mechanisms in vascular plants, which would have emerged early in the evolution of plants (Hodge, 2004). Mycorrhizal association did not detectably influence fork or link development in our study.

### 3.4 Investment in Root Biomass and the Functional Balance (allometric growth) Mechanism

Most of a root’s enzymatic activity and nutrient absorption occurs in a short hairy zone of low suberisation generally close to growing tips, and its development is an important physiological component of fitness. Interestingly, we observed a well-developed hairy region extending throughout the root length (Fig. 6A), which highlights the critical importance of maximizing the nutrient acquisition function in our nutrient-limited environment. Root surface area emerged as the lead biomass investment correlate (Fig. 1), consistent with its established role as the belowground analogue of leaf area in resource acquisition (Jackson et al., 1997). As expected from geometric principles, the thinnest roots (0–0.6 mm diameter class) contributed the greatest total surface area across all substrates but rhyolite (Fig. 6B). Generally, for a given biomass fine roots maximize absorptive area per unit carbon invested (i.e. fine roots are C-cheap), which is a key tradeoff of the root economics spectrum (Freschet et al., 2021). In rhyolite plants had the largest surface area for the whole root system, but this was mostly represented by the medium-thickness class (0.3–1.2 mm; Fig. 6B), which is consistent with roots generally being more voluminous (Fig. s6B). Conversely, in schist plants exhibited the smallest surface areas for the thinnest class (and the total root system; Fig. 6B. Together with granite, schist plants also supported largest segment surface area (Fig. 6E), which was primarily driven by greater segment length (Spearman’s *r* = 0.89, *p* < 0.01). This suggests that in schist and granite plants used the scarce resources to extend individual root segments rather than proliferating shorter, finer laterals. Notably, plants in basalt showed comparatively larger total surface areas within the thinnest diameter class (0–0.3 mm), but had low segment surface area (Figs. 6E), which were numerically abundant and had high tips number (5B, 3A and s4D). This configuration is consistent with an intensive exploration strategy — maximizing soil contact points at minimal individual segment cost — as opposed to the length-based exploration strategy observed in granite and schist.

**Fig. 6.**
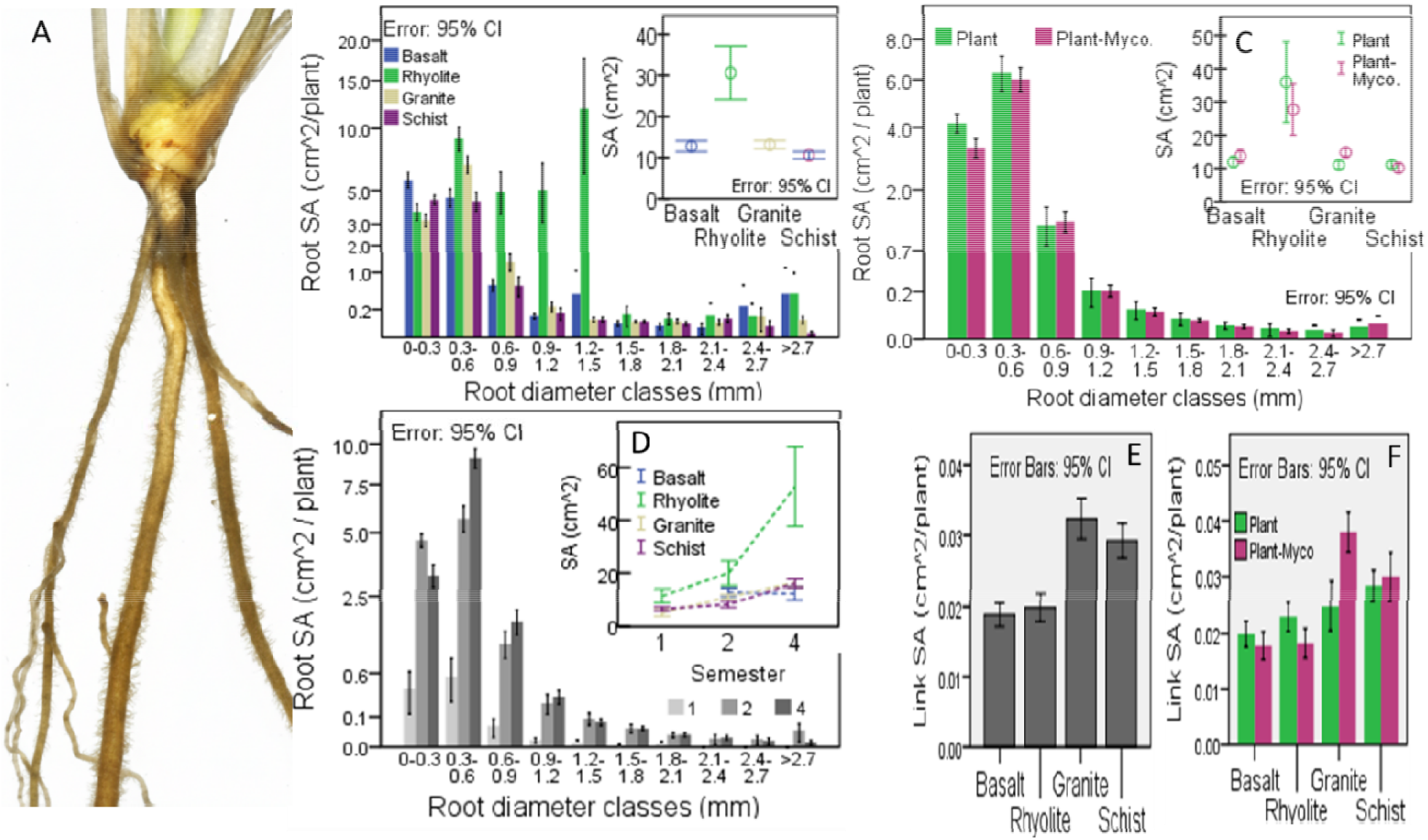
Root and link surface areas. (A) Evidence of root hairs throughout the root length in a mycorrhized plant grown in rhyolite for 129 days. (B) Rock, (C) mycorrhiza, and (D) time effects on root surface area across thickness classes and the whole root system (inset). Root segments surface area variation with (E) rock and (F) mycorrhiza. Values represent averages.

Over time, the sharpest surface area increment occurred in rhyolite (Fig. 6D inset), implying that this substrate provided the most favorable geochemical conditions to sustain continued investment in absorptive tissue, permitting sustained secondary development of fine roots. This root class is energetically more expensive to maintain, as shown by Pregitzer et al. (2002) in a N investment per root class study. Results are also consistent with the observed thicker roots and faster, exponential growth in this rock (Fig. 6B). To lesser extent, plants in schist and granite also expressed increased root surface area over time, while in basalt growth plateaued after eight months of moderate expansion— a pattern consistent with resource exhaustion after initial abundance (Fig. 6D inset). Overall, the finest roots (0– 0.3 mm) contributed to surface area expansion only during the first two growth semesters, whereas the medium-diameter class (0.3–0.9 mm) continued to accumulate surface area throughout the entire growth period (Fig. 6D). This ontogenetic shift reflects a documented allometric resource redistribution from finest to thicker roots as plants matured, diminishing investment in absorptive surface proliferation, and consolidating the existing resource transport (and structural support) pathways (Zhou et al., 2022).

Interestingly, mycorrhiza inoculation reduced overall investment in the surface area of the finest roots (0 - 0.3mm; Fig. 6C), most clearly at the root segment scale in rhyolite (Fig. 6F). This is consistent with the outsourcing strategy end member (*vs*. do-it-yourself) along the collaboration gradient of the root economics space, whereby the extraradical hyphal network supplements fine root absorptive function, reducing the plant need to maintain high surface area in the exploratory root class and freeing carbon for reallocation to other structural or metabolic domains (Bergmann et al., 2020). In granite, however, inoculation slightly stimulated investment in both total and segment root surface areas, suggesting a context-dependent shift toward plant-fungus competition for nutrients — a pattern analogous to the “do-it-yourself” acquisitive strategy end member, where the carbon “subsidy” demanded by the fungus exceeds the benefit on this particular substrate (Bergmann et al., 2020).

For early vascular plants of the Devonian times (c. 400 million years ago), maximizing nutrient absorption surface, and strategizing its placement in relationship with the chemical energy map of its environment (e.g. local mineralogy, moisture gradients, and organic matter pockets) - alongside above ground light distribution was likely a critical evolutionary leap (Raven and Edwards, 2001; Kenrick and Strullu-Derrien, 2014). The progressive elaboration of this acquisitive belowground interface, oftentimes aided by far-reaching fungal networks ultimately produced root systems that in modern temperate grasslands exceed leaf area by at least an order of magnitude (Jackson et al., 1997) — a striking inversion given that leaves access carbon and solar energy with comparatively little structural investment. This belowground interface requires c. 40% of net C fixed by photosynthesis allocated to fine roots alone (Jones et al., 2009), underscoring how central root surface area is to whole-plant carbon economics. Globally, total root surface area across all terrestrial ecosystems is estimated to reach 2.0 × 10^9^ km^2^, approximately 14 times the total land surface area (Jackson et al., 1997) — a figure that places the root-soil interface among the most extensive biological surfaces on Earth, and one whose functional properties are profoundly shaped by the mineralogical heterogeneity of the explored substrates.

Below-ground C allocation is reflected in root volume, which integrates plant investment in new resource exploration and acquisition – including uptake of nutrients such as P, metal cofactors necessary for enzymatic functions – as well as transport and structural support. Rhyolite supported by far the largest total root volume, followed by granite, basalt and schist (Fig. s6A). Fine roots dominated the total volume, with a peak at 0.3-0.6mm diameter class in basalt, granite and schist, while rhyolite peaked at thicker 1.2-1.5mm (Fig. s6B). Rhyolite is a felsic, relatively more acidic volcanic rock that, despite being base-poor, promotes greater phosphate supply to the rhizosphere through weathering (Alves-Brilhante et al., 2017; Zaharescu et al., 2019). This geochemical condition, combined with sustained availability of other critical nutrients appears to have permitted greater below-ground C allocation into thicker roots and larger overall biomass (root and shoots) biomass in this substrate (Fig. s3D). A sustained phosphate supply from rhyolite weathering has been documented in this system previously (Zaharescu et al., 2019), which could have provided the chemical energy necessary for overall root buildup.

Over time, root volume increased consistently in rhyolite and granite, plateaued after the second semester in basalt, and an increased after an initial plateau in schist (Fig. s6C), indicating substrate-dependent adjustments in the rate and trajectory of belowground investment.

Mycorrhizal inoculation produced substrate-specific effects on root volume allocation (Fig. s6D-G): in rhyolite, fungi reduced the volume of thin prospective (0.3-0.9 mm) roots; in schist, fungal presence simultaneously reduced investment in thin roots while increasing volume in medium-thickness transport roots; and in basalt and granite, inoculation stimulated transport medium-thickness root volume. This substrate-contingent variability suggests an adaptive repositioning of symbiosis along the “outsourcing” to “do-it-yourself” collaboration gradient of the root economics space (Bergmann et al., 2020), extending allometric resource redistribution from the plant alone to the broader fungus-plant system. This pattern warrants further investigation across a wider range of lithologies.

Biomass volume is distributed along growth vectors, which direct limited internal energy towards areas with external resources. Across the 797 analyzed plants, the average total root length (a global growth investment indicator) was 137cm ± 96 SD (range 9.8 – 1,097cm per plant). Rhyolite supported the greatest total root length (mean 179cm), followed by basalt (134cm), schist (114cm) and granite (112cm; Fig. 7A). Plants on granite also displayed 11-26% smaller specific root length (SRL - a proxy of growth by resource investment; cm/g dry biomass) compared to other substrates (Fig. 5D), indicating a more conservative growth strategy and greater root tissue density. Of thickness classes, the fine prospective roots (0-0.3mm diameter) dominated total length in basalt and schist, where they represented about 70% of the total length; in rhyolite and granite thicker roots (0.3 – 1.2mm, mostly involved in transport; Pregitzer et al., 1997) contributed proportionally more, particularly in rhyolite (Fig. 7C). This rock-dependent length-thickness relationship, was supported by significant length correlations between thick roots (0.9 - 1.2mm diameter) and total root system in rhyolite (Spearman r = 0.44, p < 0.01), and between fine roots (0 - 0.3mm) and total root system in basalt, granite, and schist (Spearman r > 0.48, p < 0.01). Allometric analysis of developmental order revealed that generally lateral (secondary) and tertiary roots had larger total lengths than primary (embryonic) roots, with largest values for all axes in rhyolite (Fig. 7B). Notably, rhyolite also supported longer lateral than tertiary roots, whereas in schist (and with higher variability in basalt) tertiary roots exceeded lateral roots in length (Fig. 7B).

**Fig. 7.**
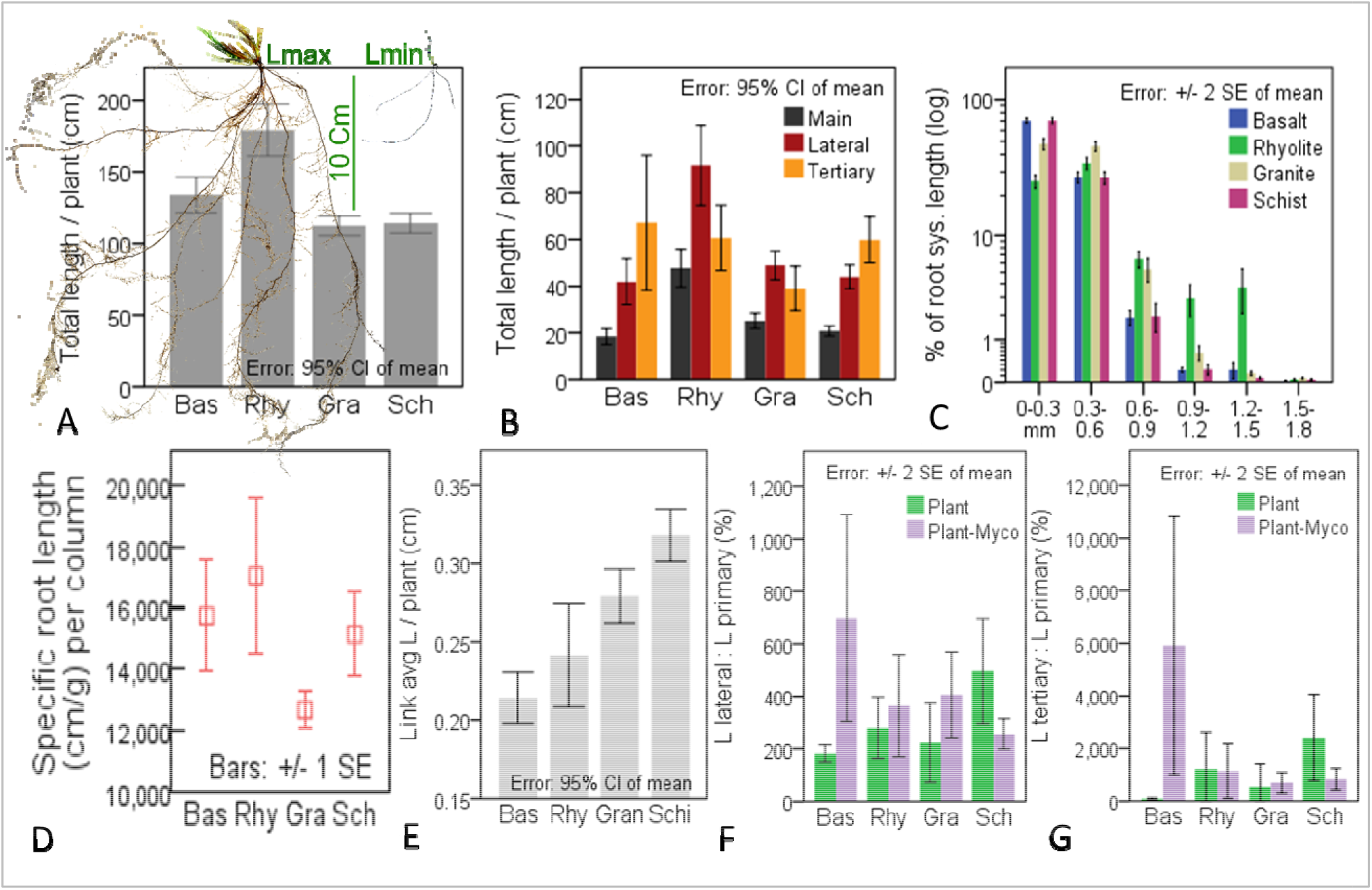
Root system and link lengths. Substrate effect on (A) root system total length, (B) total length of various root orders, (C) relative length of various root thickness classes for an average plant (calculated as % of total root system length), and (D) specific root length (total length normalized to root biomass); N=797 plants. Example of maximum and minimum root length plants in (A) are from rhyolite and schist, respectively. (E) Link average length (cm) as a function of substrate; N= 780 plants. Effect of arbuscular mycorrhiza on the Allometric Growth Index of (F) lateral to primary and (G) tertiary to primary root length ratios for an average plant; N=311 plants.

Root segments (links) – fundamental length units (between branching nodes) that intercept energy-favorable areas (Fitter and Stickland, 1991) averaged 0.2 ± 0.1cm SD (range 0 – 1.61 cm) in length, reflecting frequent root segmentation in *B. dactyloides*. The shortest segments were developed in basalt (Fig. 7E), coinciding with denser branching network (segments and axes; Fig. 5B, Fig. 4B); the longest were in schist, followed by granite and rhyolite (Fig. 7E). In rhyolite, the segment length correlated with lengths of all individual diameter classes (Spearman r > 0.38; p < 0.01), indicating proportional growth. In the other substrates, segment lengths showed negative relationship with fine roots length (<0.03mm; Spearman r < -0.33, p < 0.01), i.e., longer fine roots were composed of shorter but more abundant segments; and positive relationship with medium - thickness roots (0.3 - 0.6mm; Spearman r > 0.33, p < 0.05).

Collectively, these patterns reflect a dynamic, substrate-dependent adaptation of root architecture development (resources acquisition) to resource availability in *B. dactyloides*. On rhyolite, sustained phosphate and multi-nutrient supply (Zaharescu et al., 2019) supported investment in a robust transport system of thicker, longer roots, with embryonic (primary) and first-order lateral roots dominating total length — a pattern consistent with deeper vertical foraging at the cost of further side branching. Schist and basalt, by contrast, promoted lateral elongation across two hierarchical/fractal orders (extensive growth strategy with low investment), at the expense of embryonic and first-order lateral root elongation. In granite, the most resource-limited substrate (Zaharescu et al., 2019), plants maintained transport capacity through vertical exploration while minimising branching investment — a pattern of particularly low SRL (to preserve scarce biomass) and a more herringbone-like topology (Fitter and Stickland, 1991).

Lateral root initiation represents a plant’s metabolic decision to capitalise on localised nutrient hotspots (e.g., nitrate), redirect carbon from mother root elongation and spread locally (usually near the tip); but P deprivation at the main tip can also stimulate branching (López-Bucio et al., 2003). Energetically, lateral elongation at the cost of deeper primary rooting, usually occuring under nutrient restriction, is an efficiency strategy as it requires less biomass invested to construct and maintain (Yang et al., 2003; Lynch, 2019; Schenk and Jackson, 2002). In such conditions, using resources for lengthening is energetically favoured over morphological change. As a consequence, roots become thinner, maximising surface area per unit biomass (Zobel et al., 2007). As growth resources decline, a slow-growth, resource-conservation strategy occurs in association with low SRL (observed in granite) - characterized by increased root tissue density (carbon buildup; Comas and Eissenstat 2009; Trubat et al., 2012; Bergmann et al. 2020). In contrast, in nutrient-richer substrates (rhyolite, basalt), plants can afford to invest in both lengthing and pronounced morphological plasticity (Freschet et al., 2015).

These patterns correspond to the two ends of Fitter’s architectural continuum (Fitter and Stickland, 1991): plants in granite and schist adopted a simpler, herringbone-like strategy (longer segments, reduced branching, lower construction cost, shorter root system; Fig. 7E, D; Fig. 2) suited to exploration efficiency in low-nutrient substrates (Berntson, 1994); while plants in rhyolite and basalt adopted a more dichotomous strategy (shorter segments, frequent branching, more absorptive root tips, longer root system; Fig. 2, Fig. 3), consistent with exploiting a heterogeneous but richer resource environment (Fitter and Stickland, 1991). This interpretation is supported by studies showing positive effects of nutrient limitation and CO_2_ abundance on root segment elongation in gymnosperms and angiosperm dicots and grasses (Fitter and Stickland, 1991; Beidler et al., 2015). The decoupling of fine segment length from total root length observed here implies that the acquisitive, most metabolically active root fraction can adjust independently of the transport-dedicated fraction — a functional uncoupling that confers greater adaptive flexibility across substrates. This also implies that when nutrients become scarce, plants trade energy investment from overall root system to lengthening functionally critical foraging roots, a strategy that has been shown in other species (Zobel et al., 2007). The negative correlation between fine root length and other thickness classes in granite (Spearman r < −0.42, p < 0.01) further supports resource trade-off allocation toward fine-root elongation in the most limiting substrate, consistent with observations in other species (Zobel et al., 2007; Taylor et al., 2014; Beidler et al., 2015).

Over experimental time, there was an increase in root total length per plant, with clearest incremental growth in schist (Fig. s7A). In rhyolite, pronounced elongation occurred in the final semester; in granite, growth plateaued after the initial four months. We also observed substantial lengthening of tertiary roots over the final semester across substrates (Fig. s7B), showing energy reallocation away from the primary axis. This coincided with lengthening of roots thicker than 0.3mm over the growth period, while thinner roots decreased significantly after 8 months across substrates (Fig. 7C). Concurrently, root segments length increased significantly during the second half of growth period in rhyolite, granite and schist (Fig. 7D). Together, these temporal patterns indicate substrate-dependent slowing of root system expansion: as the initial pulse of mineral weathering subsides, plants consolidate rather than extend their root networks — investing in the elongation of existing transport pathways rather than producing new prospective segments, thereby deepening their foothold within the established herringbone domain.

Fungal symbiosis did not significantly affect total root length (not shown), but exerted substrate-specific allometric effects. In basalt, fungal presence approximately doubled tertiary root length (Fig. s8C); in schist, fungi shortened tertiary roots while lengthening primary and lateral roots (Figs. s8B, C). These contrasting responses were reflected in the allometric growth index (AGI; ratio of secondary and tertiary to primary root length), which showed increased lateral and tertiary root lengthening at the expense of the primary root in basalt, but the reverse pattern in schist (Figs. 7F, G). In rhyolite — the most nutrient-permissive substrate — fungal presence reduced total transport root length (> 0.3 mm), while in granite (a more restrictive substrate) it increased it (Figs. s8D–G).

Surprisingly, these results indicate that mycorrhizal fungi operate primarily at the allometric scale rather than the whole-system scale (a functional balance mechanism), selectively redirecting root investment in a substrate-dependent manner: stimulating lateral nutrient pathway expansion in basalt (at the cost of embryonic roots), lengthening primary pathways in schist, shortening transport pathways where nutrients are more accessible (rhyolite), and extending them where they are not (granite). This functional repositioning is consistent with the collaboration gradient of the root economics space (Bergmann et al., 2020), and suggests that the plant–fungus partnership functions as an integrated allocation system whose architecture is tuned to the local geochemical context.

The evolutionary antiquity of this substrate-responsive symbiosis is remarkable. Well-preserved fungal endosymbioses — including arbuscule-like structures, vesicles, and spores assigned to Glomeromycota and Mucoromycotina — have been documented in Rhynie Chert deposits dating to approximately 407 Ma (early Devonian; Strullu-Derrien et al., 2014). This deep evolutionary precedent suggests that substrate-specific plant–fungus allometric feedbacks were a feature of early terrestrial colonisation, and may have contributed to the successful diversification of land plants across geochemically diverse landscapes throughout the Phanerozoic, laying the foundation for the root plasticity we observe today.

Overall, the ability of plants to optimize root length parameters both, (i) when seeds fall in favorable environments (at external nutrient cost), or (ii) unfavorable environments (at internal biomass cost) is a functional balance mechanism advantageous in the variable ecosystem context. Such ability allows plants to either (i) fill substrate volume more completely (by large plastic changes), depriving other species of space even when resources are not affected (a less efficient use of internal resources for fast/extensive external resource acquisition, as reported in invasive grasses; Vaness et al., 2014); or, (ii) more efficiently use aerially-derived carbon to counterbalance the lack of other nutrients (more restrictive plastic changes such as less branching, thinning and lengthening of segments and increasing their tissue density) and increase the exploitation efficiency (Zobel et al., 2007; Berntson, 1994). This is a root-scale response to the Functional Equilibrium Hypothesis (Brouwer, 1963; Freschet et al., 2015), which postulates that at the plant scale, limitations in one resource determines the plant to invest in the organ responsible for obtaining that resource; thus linking (through plasticity) the phenotypic response to one resource availability (e.g. belowground nutrients) with the response to the availability in another resource (e.g. aboveground light).

### 3.5 Discussion of Rock and Symbiosis Effect on Geometric Choices

*B. dactyloides* consistently operated within the herringbone architectural domain across all four substrates, yet expressed mineralogically-dependent variation within that domain, from near-dichotomous branching (intensive exploitation) in basalt to nearly pure herringbone (extensive exploration) in granite. This central finding indicates that mineralogy did not simply modulate growth rate, but it determined how the root system allocated finite mineral-derived nutrients and air-acquired carbon between exploration and exploitation, branching and elongation, and self-construction versus symbiotic outsourcing. This is a strong expression of substrate influence on species-level growth strategy, and is manifested as food foraging performance differences among substrates spanning from increased exploitation efficiency per unit tissue invested (captured by size-independent traits including segment elongation and reduced tip production) in granite and schist, to increase exploration potential at the cost of efficiency (size-dependent traits) in basalt and rhyolite (Berntson, 1994).

A synthesis of substrate mineralogy’s role in root developmental strategy is displayed in Fig. 8. Among nutrient-richer substrates, mafic basalt stimulated the densest branching, shortest segments, and highest tip density, which is an architecture optimized for thorough volumetric contact and intensive acquisition of diffusion-limited resources such as P and Fe (Fitter, 1987; Dunbabin et al., 2004). Felsic rhyolite, sustained by weathering-derived phosphate (Zaharescu et al., 2019), supported a qualitatively different response — with longer, thicker transport roots growing preferentially along embryonic and first-order lateral axes — consistent with the greater morphological plasticity and root length investment per unit biomass that nutrient-rich environments are known to permit (Freschet et al., 2015). This represents a shift from exploration toward exploitation and pathway consolidation (Lynch, 2019; Pregitzer et al., 2002). At the nutrient-poor end, granite drove development towards the established herringbone phenotype: minimum branching, low SRL, long segments and primary investment in preserving biomass over developing complexity, maximizing soil volume explored per unit tissue at minimum construction cost, aligning with Fitter and Stickland (1991) and Berntson (1994). Schist, the metamorphic substrate determined the highest root tissue density, and tertiary spreading dominating hierarchical elongation, conferring exploitation efficiency over exploration breadth.

**Fig. 8.**
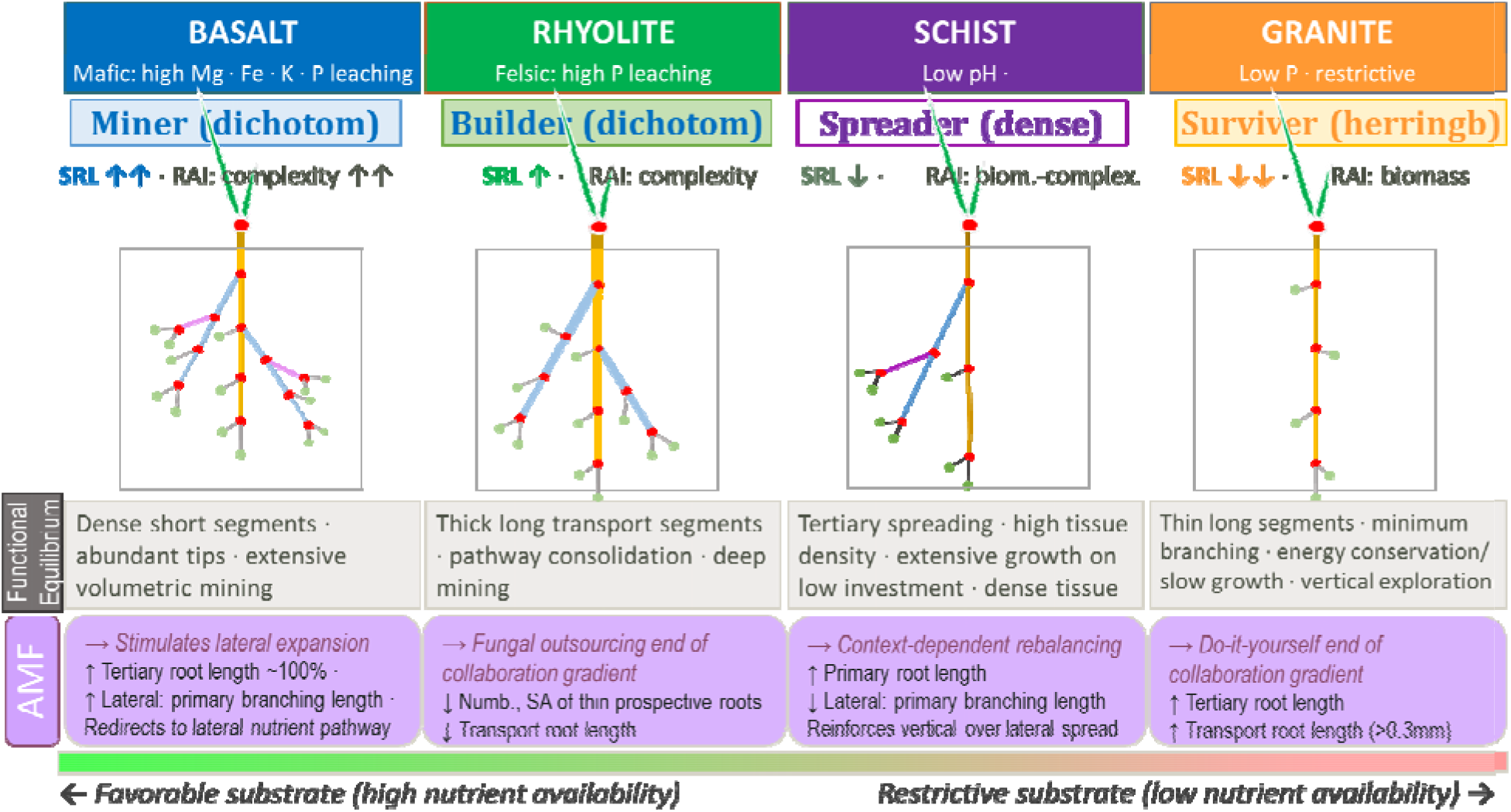
Root geometric strategy across mineral substrates. Conceptual diagram of B. dactyloides root geometry underlying changes in B. dactyloides root geometry as affected by four mineral substrates and presence of arbuscular mycorhyza, ordered along a nutrient availability gradient. Substrate chemistry drives a shift from herringbone (sparse branching, long segments, biomass-conservative) to dichotomous-like topology (dense branching, short segments, high complexity). SRL = Specific Root Length; RAI = Root Allometric Index.

Temporal consolidation across all four substrates after an initial growth phase characterized root development. This manifested as progressive lengthening of transport segments displacing fine root proliferation after half year of growth, which reflects a substrate-wide ontogenetic shift from exploration (energetically costly) to reinforcement as the initial weathering nutrient pulse subsided (Giehl et al. 2014; Pregitzer et al., 2002). Convergence of some root architectural responses to nutrient availability (e.g. segment elongation under nutrient dilution) was documented across gymnosperms (*Pinus taeda*; Beidler et al., 2015) and angiosperms (dicots and grasses; Fitter and Stickland, 1991), supporting the evolutionary conservation of important trait–substrate feedbacks across vascular plant lineages.

Mycorrhizal symbiosis acted at the allometric rather than whole-system scale, redirecting growth of specific root classes in a substrate-specific pattern: amplifying lateral branching in basalt, extending and increasing density of primary pathways in schist, shortening transport roots in nutrient-permissive rhyolite, and lengthening them in restricting granite. This rebalancing is consistent with the collaboration gradient (*vs*. conservation) of the root economics space (Bergmann et al., 2020; but see Bunn et al., 2024), integrating the plant–fungus system into a holistic nutrient allocation unit, where the small diameter fungal hyphae use surplus carbon (and reduced plant sink demand due to low nutrient) to substitute for much thicker, energetically costly fine root construction where nutrient access is inadequate, and supplement transport capacity where nutrient is more easily accessed (Hodge, 2004; Bunn et al., 2024). The breadth of geometric plasticity documented here within a single grass–fungus partnership likely recapitulates selection pressures operating since the earliest plant colonization of mineralogically heterogeneous land surfaces (Kenrick & Strullu-Derrien, 2014).

## 4. Conclusion

When germinated and grown in four contrasting mineral substrates, *Bouteloua dactyloides* consistently reallocated photosynthetically fixed carbon from above-ground organs toward the root system, prioritizing survival and nutrient acquisition over shoot expansion in the absence of organic soil amendment. Plants deployed a suite of substrate-specific architectural strategies to maximize fitness over the two-year experiment. Traits associated with exploratory effort were decoupled from those associated with biomass construction in a rock-dependent manner, indicating that substrate determines not only how much plants invest belowground, but how that investment is deployed.

Root topology organized along a wide region of the herringbone–dichotomous domain (of the herringbone – dichotomous continuum), from the biomass-conservative, low-branching herringbone strategy of nutrient-poor substrates to the complexity-intensive, densely branched near-dichotomous architecture of nutrient-rich substrates. Across substrates, the root system operated as an allometrically integrated energy allocation unit, partitioning limited rock-derived resources and photosynthates between exploration and exploitation, branching and elongation, and fine root proliferation and transport consolidation, in ways that were predictable from substrate mineralogy.

As mineral -derived nutrients declined over time, plants progressively shifted from prospective fine root proliferation toward reinforcement of established transport pathways, which is an ontogenetic consolidation consistent with the Functional Equilibrium Hypothesis operating at the architectural rather than whole-organ scale.

Mycorrhizal symbiosis extended this functional balance from the plant to the broader plant–fungus system, redirecting investment among the different root orders, rather than altering the root system as a whole. This substrate-specific allometric effect implies that in the root economics space, the plant-fungal collaboration is not a fixed, species-level trait, but a dynamic architectural response to local geochemistry, with the plant–fungus partnership functioning as an integrated allocation unit whose geometry is determined by the mineral substrate it inhabits.

The breadth of phenotypic plasticity documented here within a single grass–fungus partnership, the convergence of geometric root responses to nutrient limitation across gymnosperms and angiosperms, and the deep evolutionary antiquity of arbuscular mycorrhizal symbiosis in early land plants together support the interpretation that substrate-tuned allometric feedbacks are evolutionarily conserved features of the vascular plant root system. These traits would have been available to natural selection from the earliest colonization of the mineralogically heterogeneous Phanerozoic land surface, and they continue to govern the architectural diversity and ecological success of grasses across the geochemically variable terrestrial environments they dominate today.

## Acknowledgements

This research was funded by National Science Foundation (NSF) grant EAR-1023215 “ETBC: Plant-microbe-mineral interaction as a driver for rock weathering and chemical denudation”. We also acknowledge additional support from NSF EAR-0724958 and EAR-1331408 grants that support the Catalina–Jemez Critical Zone Observatory (CZO), the Biosphere 2 REU program, NSF EAR-1263251 and NSF EAR-1004353 (http://www.b2science.org/outreach/reu), United States-Mexico Commission for Educational and Cultural Exchange (COMEXUS): the Fulbright-Garcia Robles Scholarship program; and Thomas R. Brown Foundation endowment to University of Arizona.

We are deeply thankful to Julie Neilson, Claudia Quilesfogel-Esparza; Emily E. Gaddis; Elise N. Munoz; Shana Sandhaus; Maria A. Palacios-Menendez; Maria O. Vaquera-Ibarra; Edward A. Hunt; Nicolas Perdrial, Julia Perdrial, Nate Abramson, Juliana Gil Loiaza, Jake Kelly, Vanessa Yubeta, Lauren Guthridge, Mathew Clark, James Olmid, Guillermo Molano, Andrew Toriello, Nicolas Sertillanges, Arturo Jacobo, Yadi Wang, Joost Van Haren, Peter Troch, John Adams and Biosphere-2 team, and the multiple other collaborators for their field, lab and theoretical contribution.

## METHODS

### Construction of Model Ecosystem

A model ecosystem experiment controlling for substrate type, biota, water and atmospheric inputs was designed, built and set up in the Desert Biome of Biosphere-2, University of Arizona (Arizona, U.S.A.) (Fig. s1). The experiment location received evenly distributed solar light, with environmentally controlled temperature and humidity in the biome, with a mean temperature of 19±4°C, relative humidity of 48±19%, and natural O2/CO2 saturation conditions. The mesocosm design consisted of **(i)** six identical chambers (experimental modules), capable of maintaining an ultraclean multiannual experiment; **(ii)** an air purification module delivering 2X HEPA filtered and 2X UV-sterilized air at equal air flow among chambers (1L air per second per experimental module; Germguardian, AC4850CAPT Digital 3-in-1 Hepa Air Purifier System); and **(iii)**, a watering system to dispense controlled volumes of sterile milli-Q water (Fig. s1; Zaharescu et al., 2017). Each experimental module (170 cm length × 0.77 cm width × 0.98 cm height) was constructed from transparent acrylic, and consisted, top-down of: **(i)** an aboveground-simulating compartment provided with air and water inputs; **(ii)** a thermally and light-protected underground-simulating compartment hosting experimental tubes; and **(iii)** a sampling compartment for collecting porewater and holding the weight of the rock-filled tubes. Experimental tubes (30 x 5 cm internal diameter), protruding from the lower to the upper compartments were tightly fit in the upper compartment using bi-color (yellow/black) silicone rings acting as gas, thermal, and light barriers. The aerial compartment experienced solar radiative heating of about 5°C above the Biome average during sunny days. Potential edge effect on the experimental environment was minimized by using a hexagonal chamber geometry with tilted walls. The experimental tubes were sealed at the lower end with a tailor-made conical acrylic cap, and provided with a nylon sample extraction luer and rubber cap.

### Treatment Design

For this study, a total of 96 tubes hosted experimental treatments representing **(i)** 4 rock substrates (basalt, rhyolite, granite, and schist), **(ii)** 2 biotic treatments (plant-microbes and plant-microbes-mycorrhiza), **(iii)** 3 treatment replicates, and **(iv)** 4 time replicates.

Tubes were filled with one of the four granular rocks, and seeded with either a plant-microbial or plant microbial-mycorrhiza mixture. The filling material had a weight (g), density (g*cm^-3^), and porosity (g water/g rock; to saturation) of: 690 (± 6), 1.32 (± 0.03), and 0.33 (± 0.007) for basalt; 690 (± 10), 1.24 (± 0.02), and 0.35 (± 0.01) for rhyolite; 640 (± 8), 1.34 (± 0.003), and 0.32 (± 0.008) for granite; and, 480 (± 5), 1.04 (± 0.02), and 0.43 (± 0.01) for schist.

### Rock Substrates Selection and Processing

Rocks used for plant growth were collected from: Meriam crater, Flagstaff, Arizona (cinder basalt), Valles Caldera National Preserve, New Mexico (rhyolite), and Santa Catalina Mountains, Arizona, U.S.A. (medium-grain Oracle granite and micaceous Pinal schist). The rock substrates were selected to cover Earth’s major unweathered crustal materials, i.e. mafic (basalt), felsic (granite and rhyolite), and metamorphic. They also differ in terms of texture, mineralogy and chemical composition, which are expected to influence weathering and plant development. A detailed description of the used materials is presented in Burghelea et al., 2015, Zaharescu et al., 2019.

Except for basalt (a tephritic deposit industrially crushed at the mining site), the rocks were processed manually by removing the weathered surfaces using a pneumatic hammer and tungsten carbide-tip chisel, crushed using jaw and roll crushers, dry- and wet-sieved to 250-500μm grain size. To further remove impurities and ions input from grinding and washing, the material was passed on a Wilfley water table, rinsed several times with 18MΩ ultrapure water, and fast dried at 85°C to limit water-rock interactions. Before the experiment, the material was 3x autoclaved (121◦C for 2h) during three consecutive days to inactivate microbes, and loaded into the UV-C sterilized (30 minutes) plexiglass columns under a laminar-flow hood provided with HEPA and UV-C air purification.

### Reconstruction of Simplified Ecosystem Association

Buffalo grass (*Bouteloua dactyloides)-* a species tolerant to low nutrient environments (Solis-Dominguez et al. 2012), *Rhizophagus irregularis* (formerly *Glomus intraradices*) arbuscular mycorrhiza (AM) fungus commonly found in found in grass plant associations (Eskandari and Danesh 2010), and native rock microbial consortia were used to reconstruct a simplified ecosystem association for this study, as follows:

Microbial inoculation was performed for all rock substrates using a natural microbial consortium extracted from fresh cinder basalt (Merriam crater, Flagstaff, AZ). The protocol was previously described in Burghelea et al., 2015. Briefly, sand-size material (<2mm diameter) was collected from pristine deposits at the top of a side eruption (35°20’3.23”N; 111°16’45.48”W), transported to the laboratory on ice in sterile bags, and preserved at 4◦C until microbial extraction. In the laboratory, 1g basalt was mixed with 95mL ultrapure, sterile water (to prevent chemical and microbial contamination), vortexed for 2min and bath sonicated (40Hz) for 2min to separate aggregates and microbes, and decanted. The supernatant was used as inoculum. Heterotrophic microbial abundance was evaluated using colony-forming microbes (CFM) counts after incubation on fungicide (200mg/L cycloheximide)-spiked R2A agar plates at room temperature. The inoculum, containing 1.43 × 10^5^ CFM mL^−1^ was further sieved through a sterile 25μm sieve to remove potential native mycorrhizae spores. A total of 90ml inoculum was mixed with the substrate in each treatment used in this study.

Seeds of *Bouteloua dactyloides* (purchased from Western Native Seed, Colorado, USA) were dehusked under optical microscope, surface-sterilized to remove potential pathogens, and pregerminated before introducing them into the wet substrate. Sterilization was performed stepwise: 95% ethanol (5min), ultrapure autoclaved water rinse, 2% sodium hypochlorite (5min), ultrapure autoclaved water rinse (3x), 0.1% sodium thiosulfate (3min), ultrapure autoclaved water (3x) (Burghelea et al., 2015). Seed sterilization was checked by incubating seeds on R2A agar plates.

This ensured maximum seed survival rates in the experiment, sterile seeds were pregerminated in Petri dishes using sterile ultrapure water. A total of 20 pregerminated seeds were added to each experimental column at 2cm depth.

Sterile spores of *Rhizophagus irregularis* mycorrhizal fungus, purchased from Premier Tech Biotechnologies, CA were suspended in ultrapure sterile water and added together with the seeds to the rock substrate (about 400 spores per column) to maximize symbiosis establishment.

Prepared columns were placed in each of the six pre-sterilized (UV-C for 30 minutes) modules in a randomized configuration to prevent edge effect on plant growth. Measured volumes of sterile ultrapure water was added manually through the outside ports to each column using a sterile 140mL polypropylene syringe, and distributed on column surfaces using an in-house designed dispenser that prevented preferential flow (Fig. s1-E). Every 2 weeks, about 100-120mL sterile ultrapure water (18MΩ) was added to each column (to near saturation), to maintain an active plant – microbiota – weathering system, and the volume was recorded for mass-balance calculations.

### Plant collection

During the experiment, planted column replicates were extracted (sacrificed) over 4 time points at 132, 252, 465 and 584 days to evaluate the temporal changes in root growth patterns. Under a laminar-flow hood, the contents of the columns were emptied into clean polypropylene trays, and plants were gently separated from the granular substrate by hand to prevent root damage. After submerging in ultrapure water to remove adhered rock particles, plants were stretched on a dark surface and photographed next to a ruler to measure above- and below-ground organ lengths.

### Root scanning and architecture analysis

A total of 797 plants were submerged in shallow water in a transparent acrylic tray, and their roots were spread out individually to minimize root segment crossings. Image acquisition was performed using an Epson Expression 10000XL optical scanner (a dual lighting system that produces shadow-free images) with a scanning area of 30×42cm. Uncompressed TIF images (300, 400 or 600 dpi pixel density) were collected, and analyzed for each plant with WinRHIZO Pro v. 2009 (Regent Instruments Inc., Canada).

Several architectural proxies were used to capture fine differences among root growth patterns, and their effect on fitness: **(i)** geometry (frequency, lengths, area, volume), **(ii)** topology (branching patterns), and **(iii)** biomass. The scanning, parametrization and data analysis workflow is presented in Fig. s2-A., with an explanatory list of measured variables in Fig., s2-B.

Altitude Topological Index (ATI), an estimation of branching patterns ranging from herringbone - like (values closer to 1) to dichotomous (values closer to 0), was inferred using the relationship between log10 of altitude (a, the largest path for nutrients by segment number) and log10 of tips number (μ, the most metabolically active and the prospective part of the root): ATI = log (a)/log(μ) (Fitter, 1987; Beidler et al., 2015). We chose ATI because this index discriminated well among rock types.

### Mycorrhizal evaluation after column extraction

After scanning, root fragments were subsampled for mycorrhiza infection rate determination, the plants were individually dried at 70 °C for 3 days, separated into above and below ground biomass, and weighed.

To determine the mycorrhization rate, 1cm root subsamples were randomly collected along the roots form an experimental column, cleared by boiling (90◦C for 15min) in 10% KOH, the solution neutralized (room temperature for 2min) with 10% HCl, followed by hot root staining (90◦C for 10min) using 0.05% trypan blue. The % rate of root mycorrhization was determined under optical microscope, by recording the number of segments showing any infection out of a total of 30 random segments (detailed in Burghelea et al., 2015, after Phillips and Hayman 1970).

### Statistical analysis

Data analysis is summarized in Fig. s2-A. Principal component analysis was used to find similarities among root variables, and to group variables into a small number of higher-order factors (principal components) that could be used as broad root development traits to explain root allometric growth. A Varimax rotation method with Kaiser Normalization was used to maximize the principal component explained variance (the fit between principal component axes and variables).

To test whether there is a variable and consistent investment in allometric growth in plants growing in different rock substrates, we developped a Root Allometry Investment Index (RAI), which compares the effort invested in complexity (regression factor scores of PC1) to effort invested in unit biomass (regression factor scores of PC2) using the formula RAI=(PC1+1)/(PC2+1) with values below 1 representing investment in biomass (PC2), and values above 1 denoting investment in complexity (PC1).

