## Supplementary Information for "Mineral geochemistry and mycorrhizal allocation define root architectural strategy during early vascular plant colonization"

### Experimental Setup

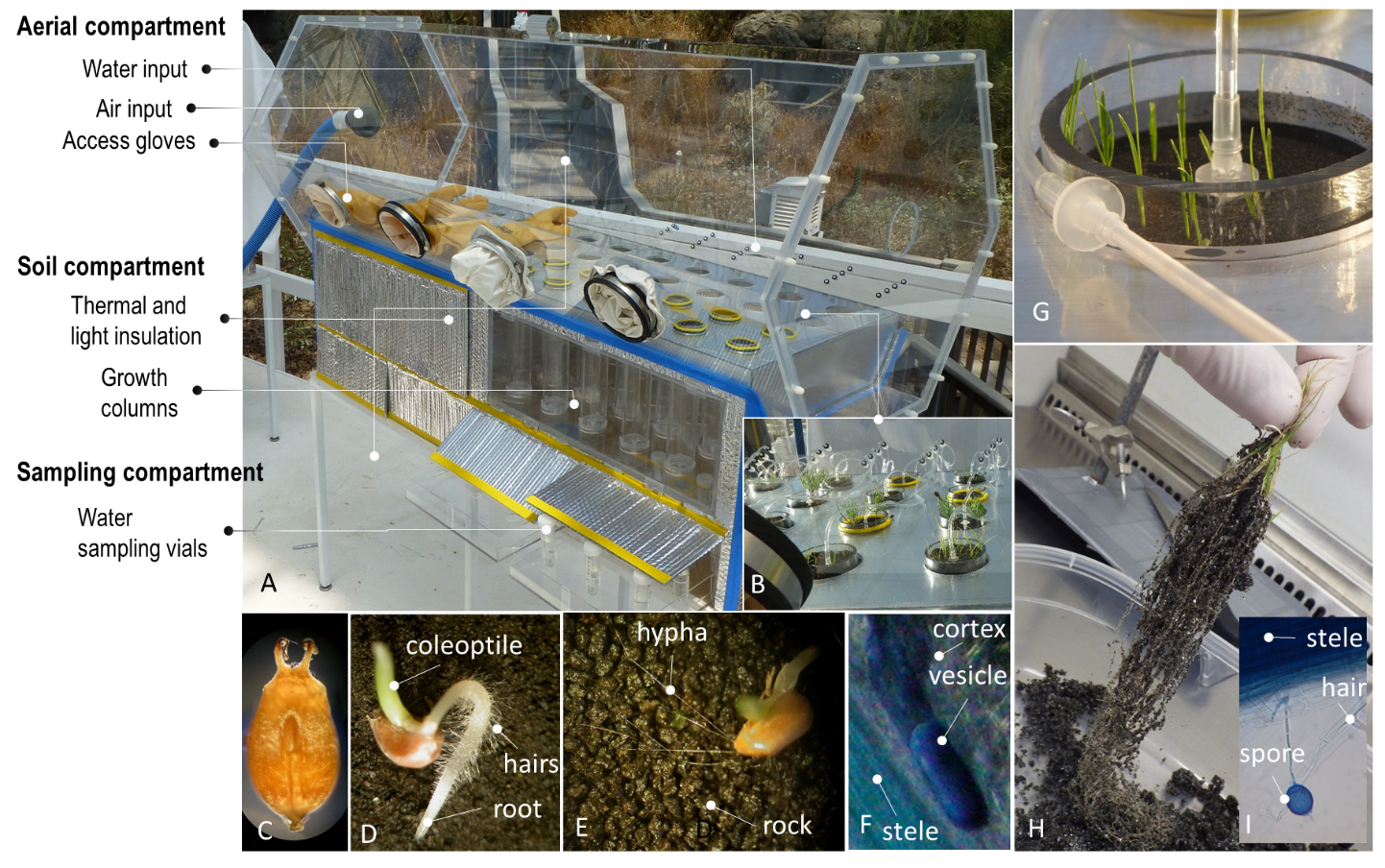

**Fig. s1 Experimental setup.** (A, B) One of the six identical growth chambers built for the experiment showing above and below ground, and sampling compartments; (C, D) Seed and germinated seed of *Bouteloua dactyloides* (buffalo grass) used in this study; (E) Optical microscope image of germinated seed and arbuscular mycorrhiza *Rhizophagus irregularisarbuscular* (VAM) hyphae connecting emerging roots with rock grains; (F) Stained root tissue showing a VAM vesicle embedded in cortical cells; (G) Plants in basalt the first semester of growth. (H) Roots extracted from schist at the end of the experiment, and adhered rock grains; (I) VAM spores emerging from plant root cortical region.

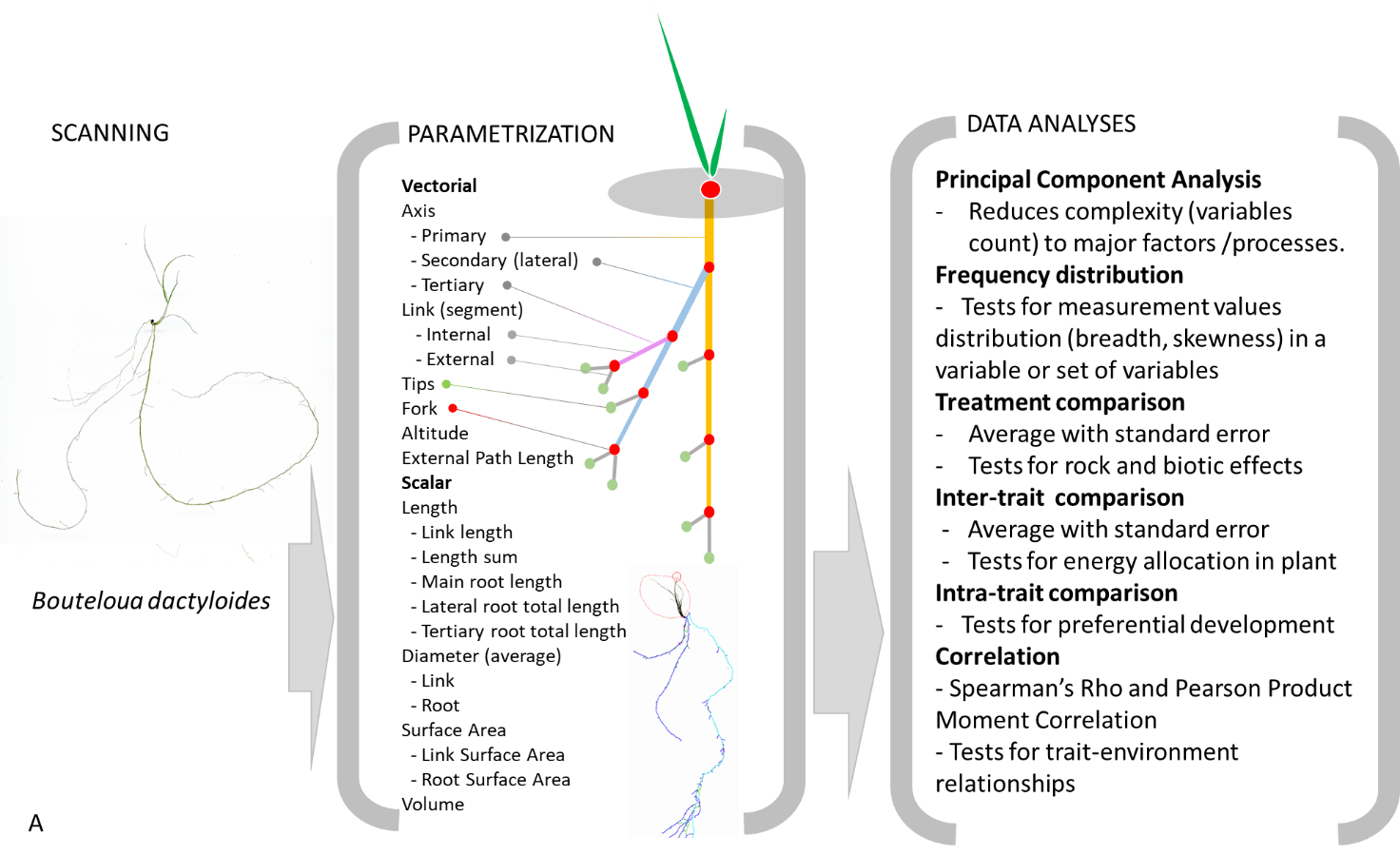

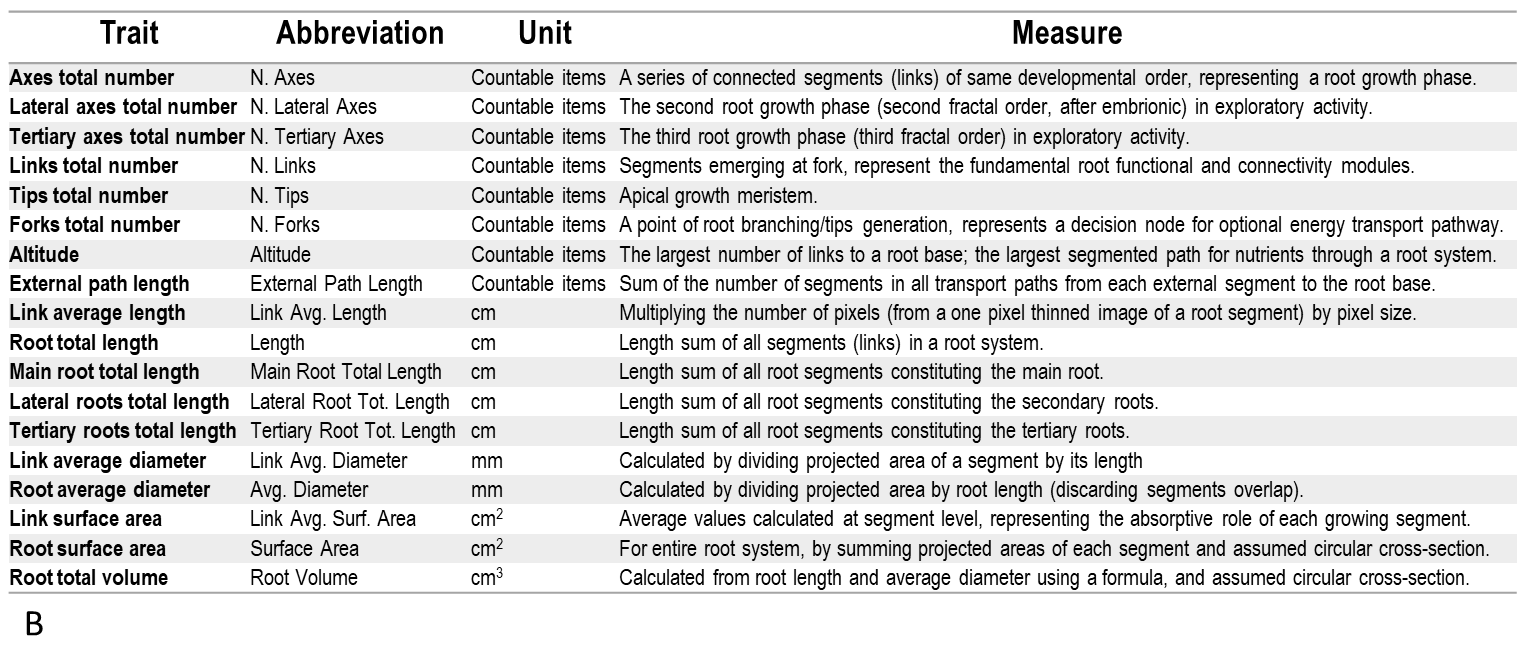

**Fig. s2 Deconstructing the root architecture and statistical logic.** (A) Description of procedural steps for the analysis of 780 *Bouteloua dactyloides* roots, together with data analysis steps that were employed.

(B) Description of measured root parameters that were relevant for statistical analysis (onthology after Régent Instruments, 2021, Lobet et al., 2015, Beidler et al., 2015).

### Major Root Characteristics

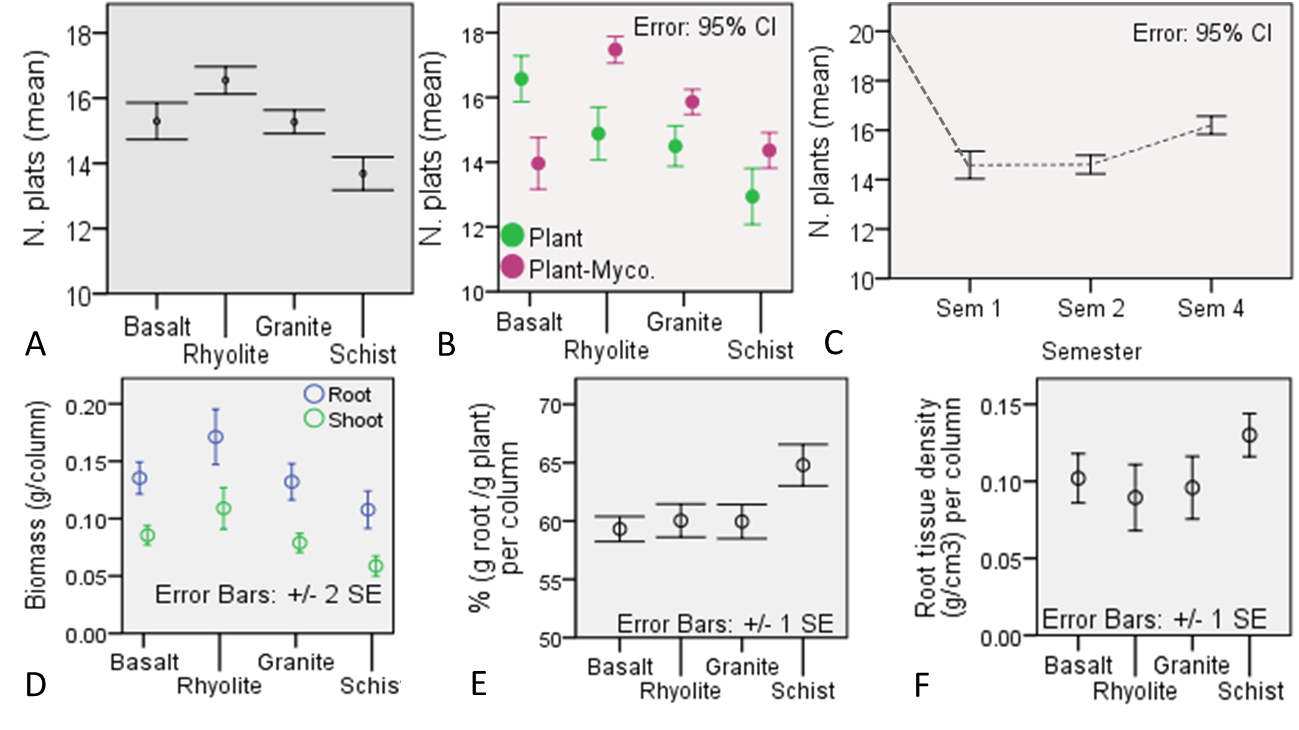

**Fig. s3 Major plant descriptors.** Survival rates of *B. dactyloides* seedlings in different substrates (A), inoculated and not-inoculated with arbuscular mycorrhiza (B) and at different sacrifice times (i.e. three month intervals; C), out of a total of 20 initial pre-germinated seeds per growth column. N = 780 surviving plants were scanned for root analysis. Plant total column biomass (D) and root mass fraction (E) pooled across the four sacrifice points and two biotic treatments, together with (F) root tissue density.

### Investment in Root Complexity

We tested the idea that global topology parameters, such as altitude and external path length, provide a refined description of root branching strategies. Altitude, a proxy for the largest sequential (exploratory) growth of root functional modules (links), was highest in schist, with an average value of 86.9 segments, and smallest in rhyolite (average altitude = 74.9 segments; Fig. s4A). This difference indicates that plants grown in schist substrate developed the largest path for nutrients (by segment number) through the root system. Altitude grew from seed to semester 1 and 2 (6 months) after which it stabilized (Fig. s4B). Mycorrhizal presence significantly diminished root sequential growth in rhyolite (Fig. s4C) likely by unloading some of the host’s exploration investment in its extended hyphal network. External path length, a measure of the entire root system density, was smaller in granite than in rhyolite and basalt (Fig. s4D), reflecting low C investment in this trait. Marked differences in root system density among rocks only occured in the last semester of the study period (Fig. s4E).

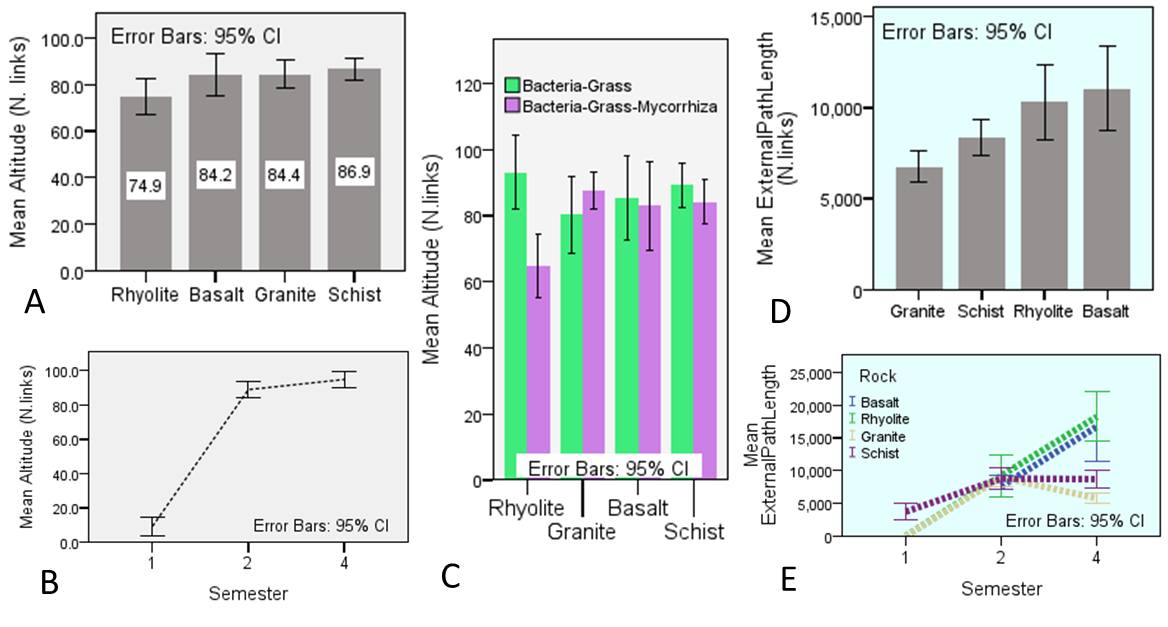

**Fig. s4 Maximum nutrient path and root system density.** Effect of (A) rock, (B) time, and (C) biota on altitude (maximum root nutrient path) in *Bouteloua dactyloides* over the 2 years experiment. External path length (root system density) as affected by (D) rocks and (E) time. N= 718 plants.

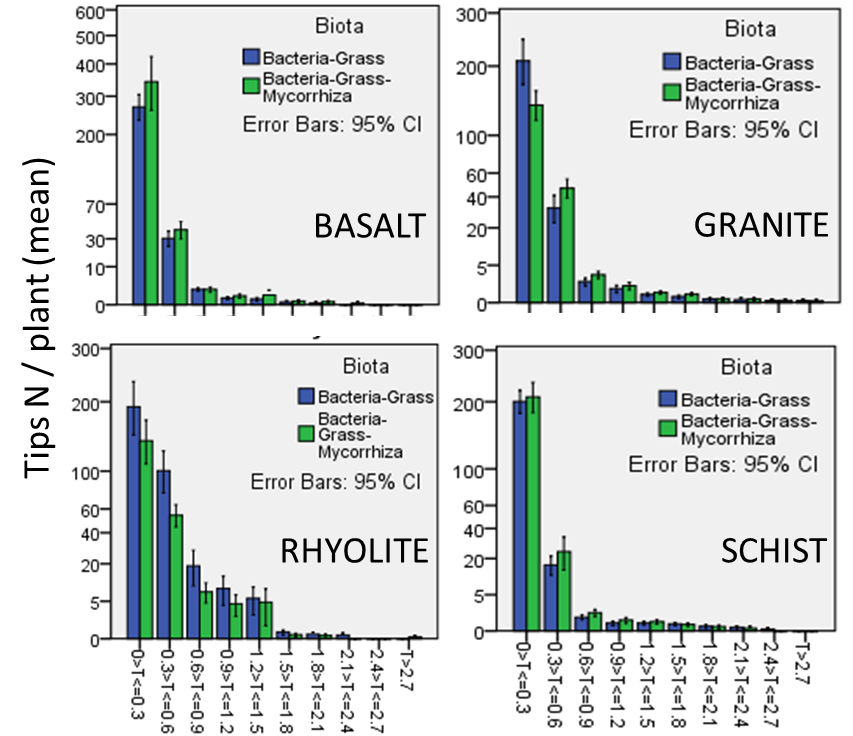

**Fig. s5 Mycorrhiza effect on root tips development.** (A) Tips abundance for each root thickness class in plants inoculated and non-inoculated with arbuscular mycorrhiza *Rhizophagus irregularis,* grown for two years in the four rock substrates.

### Investment in Root Biomass

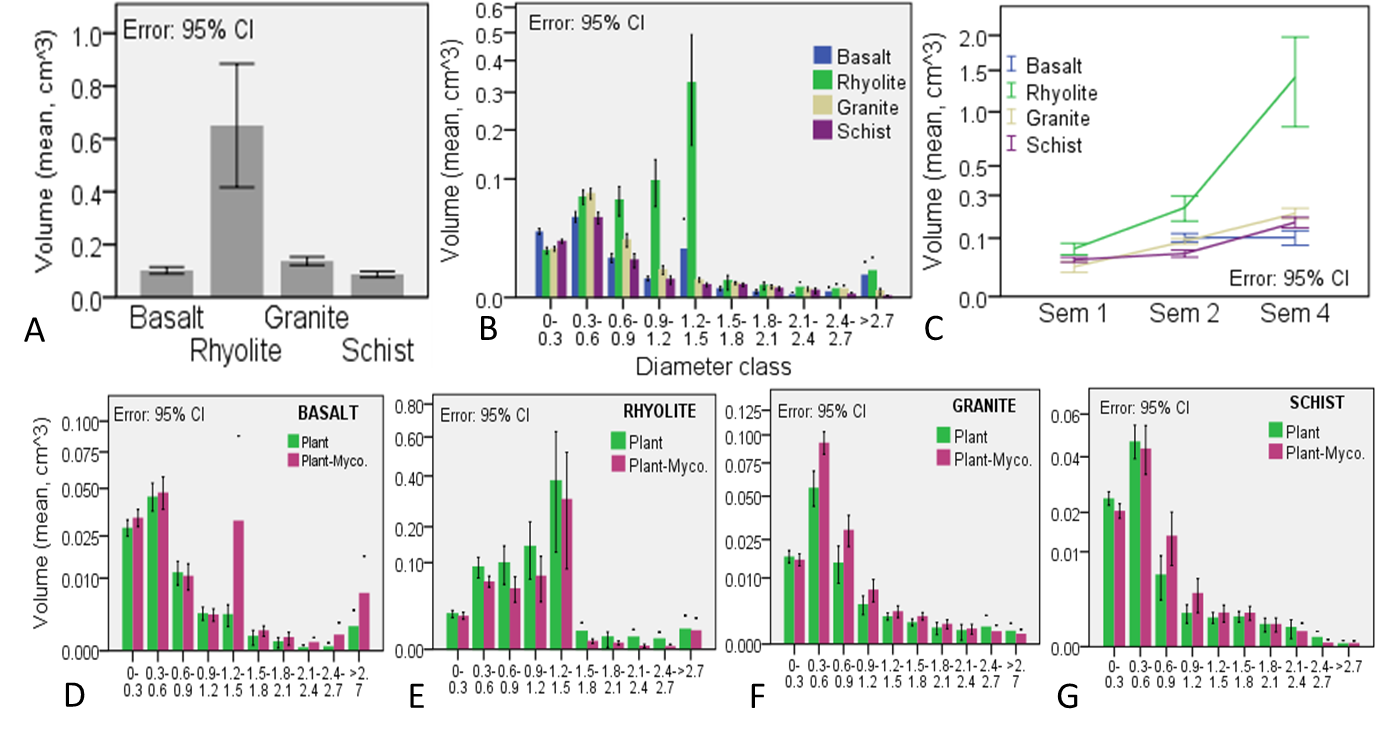

**Fig. s6 Root volume.** Rock effect on (A) total root volume, and (B) root volume by diameter class. (C) Root volume over time. (D - G) Mycorrhiza effect on root volume across root thickness classes in the four rocks.

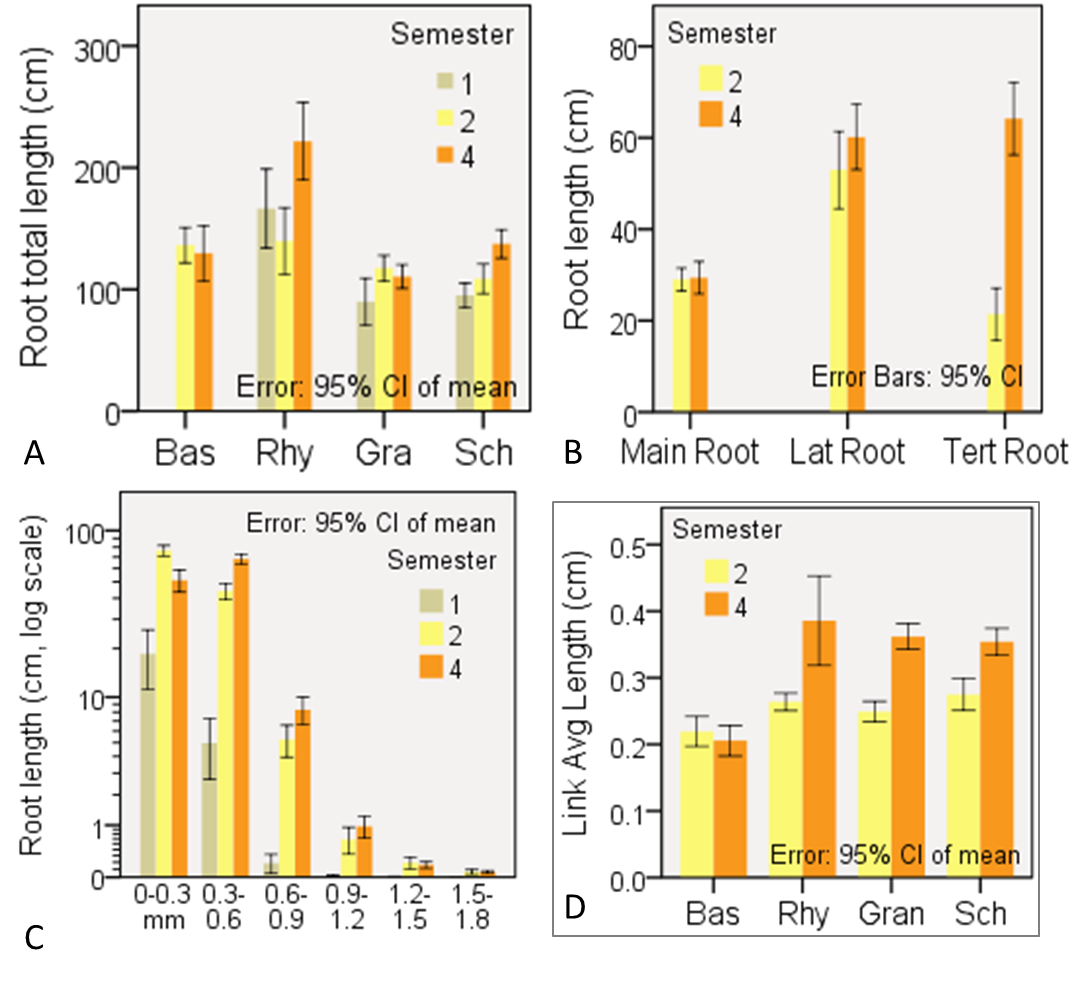

**Fig. s7 Root length over time.** (A) Root system total length (mean per plant) at different sacrificial time points in the four substrates. Time changes in mean root length (per plant) across (B) root orders, and (C) root thickness classes. (D) Link average length (per plant) changes in time in the four substrates. Each semester represents 3 months of growth.

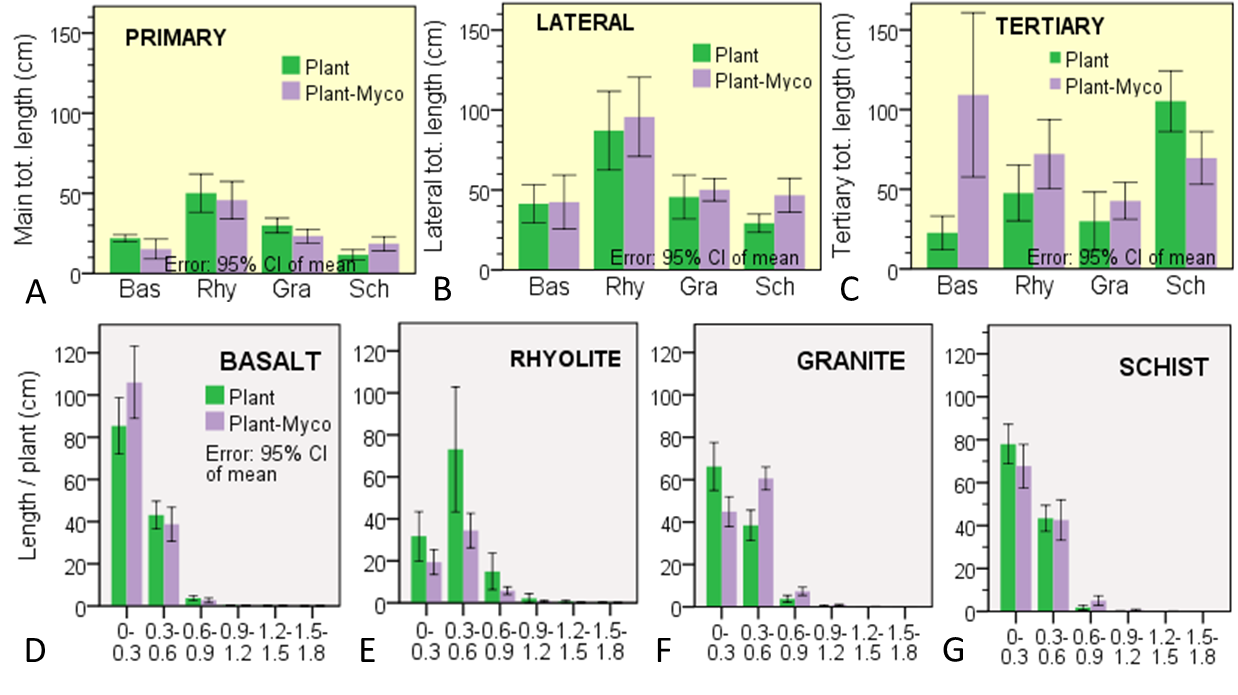

**Fig. s8 Mycorrhiza effect on root length.** Total lengths of (A) main, (B) lateral, and (C) tertiary roots as affected by arbuscular mycorrhiza in various rock substrates. (D-G) Mycorrhiza effect on total root length across thickness classes in the four rocks. Measurements were taken from the fourth semester plants.

**REFERENCES SI**

Régent Instruments Inc. (2021). WinRHIZO: Image analysis system for root measurement. User manual. Quebec, Canada. Retrieved from https://www.regentinstruments.com/assets/images_winrhizo/WinRHIZO_2021.pdf.

Lobet G, Pound MP, Diener J, Pradal C, Draye X, Godin C, Javaux M, Leitner D, Meunier F, Nacry P, Pridmore TP, Schnepf A. (2015 ) Root system markup language: toward a unified root architecture description language. Plant Physiol.167(3):617-27. doi: 10.1104/pp.114.253625. Epub 2015 Jan 22. PMID: 25614065; PMCID: PMC4348768.

Beidler KV, Taylor BN, Strand AE, Cooper ER, Schönholz M, Pritchard SG. (2015) Changes in root architecture under elevated concentrations of CO₂ and nitrogen reflect alternate soil exploration strategies. New Phytol. 205(3):1153-1163. doi: 10.1111/nph.13123. Epub 2014 Oct 28. PMID: 25348775.
